# Nucleus accumbens dopamine release reflects Bayesian inference during instrumental learning

**DOI:** 10.1101/2023.11.10.566306

**Authors:** Albert J. Qü, Lung-Hao Tai, Christopher D. Hall, Emilie M. Tu, Maria K. Eckstein, Karyna Mishchanchuk, Wan Chen Lin, Juliana B. Chase, Andrew F. MacAskill, Anne G. E. Collins, Samuel J. Gershman, Linda Wilbrecht

**Affiliations:** Department of Psychology, University of California, Berkeley, CA, 94720, USA; Center for Computational Biology, University of California, Berkeley, CA, 94720, USA; Helen Wills Neuroscience Institute, University of California, Berkeley, CA, 94720, USA; Sainsbury Wellcome Centre for Neural Circuits and Behaviour, University College London, London, W1T 4JG, UK; Google DeepMind, London, UK; Department of Neuroscience, Physiology and Pharmacology, University College London, UK; Department of Psychology and Center for Brain Science, Harvard University, Cambridge, MA, 02138, USA; Center for Brains, Minds, and Machines, Massachusetts Institute of Technology, Cambridge, MA, 02139, USA

## Abstract

Dopamine release in the nucleus accumbens has been hypothesized to signal reward prediction error, the difference between observed and predicted reward, suggesting a biological implementation for reinforcement learning. Rigorous tests of this hypothesis require assumptions about how the brain maps sensory signals to reward predictions, yet this mapping is still poorly understood. In particular, the mapping is non-trivial when sensory signals provide ambiguous information about the hidden state of the environment. Previous work using classical conditioning tasks has suggested that reward predictions are generated conditional on probabilistic beliefs about the hidden state, such that dopamine implicitly reflects these beliefs. Here we test this hypothesis in the context of an instrumental task (a two-armed bandit), where the hidden state switches repeatedly. We measured choice behavior and recorded dLight signals reflecting dopamine release in the nucleus accumbens core. Model comparison among a wide set of cognitive models based on the behavioral data favored models that used Bayesian updating of probabilistic beliefs. These same models also quantitatively matched the dopamine measurements better than non-Bayesian alternatives. We conclude that probabilistic belief computation contributes to instrumental task performance in mice and is reflected in mesolimbic dopamine signaling.

## Introduction

Reinforcement learning (RL) algorithms hypothesize that, in value based decision making tasks, animals maintain an action value map and update it using reward prediction errors (RPE), the difference between the observed and predicted reward associated with the chosen action [1]. The growing family of RL models has proved remarkably successful for explaining trial-by-trial changes in behavior [2–4] and the responses of midbrain dopamine neurons in rodents and primates [5–7]. However, these standard RL (SRL) models may not account as well for behavioral and neural data when sensory observations are not sufficient to determine optimal behavior, for example, when the underlying reward distribution changes over time due to the existence of a hidden state[8]. To solve such tasks, evidence suggests that the brain may represent a “belief state” (a probability distribution over hidden states), updated by Bayesian inference [9–14], or possibly learned implicitly in an end-to-end fashion [15]. Additionally, updates to these belief state-dependent action values may be encoded by dopamine neurons [9–13].

The significance of Bayesian inference becomes apparent when studying behavior in a two-armed bandit task (2ABT, Fig. 1A), a serial reversal learning task where rewards for correct choices are delivered on a probabilistic schedule, and the correct port switches between two available options across blocks of trials within a single session. The fact that rewards are delivered on a probabilistic schedule generates ambiguity as to which port is currently the rewarded port following an unrewarded trial. Both mice and human subjects trained in this task are capable of maintaining stable behavior within a block, and then rapidly adapting after reward contingencies reverse following a block switch [16–18]. The simplest versions of standard RL models do not capture choice stability within a block and rapid switching when the rewarded port changes [17, 18]. Bayesian inference may offer better models of behavior in tasks like the 2ABT because they capture stable within-block choice and rapid between-block switching [12]. Past studies have also shown that dopamine signaling may be better explained by models accounting for belief states [9–11, 13, 19]. However, the additional explanatory power of Bayesian models with respect to RL models is currently ambiguous, given that a growing family of complex RL models, geared with more parameters and nuanced functions, are capable of generating adaptive behaviors [18, 20]. A recent study of human behavior in the 2ABT found that a Bayesian model and an RL model variant equipped with counterfactual value updates both provided complementary explanations of behavior, but neither was decisively superior [17]. Recent behavioral analyses of mice performing the 2ABT also suggested a purely Bayesian account of reversal learning can overestimate mouse switch probability and fail to account for choice stickiness [18].

**Figure 1.**
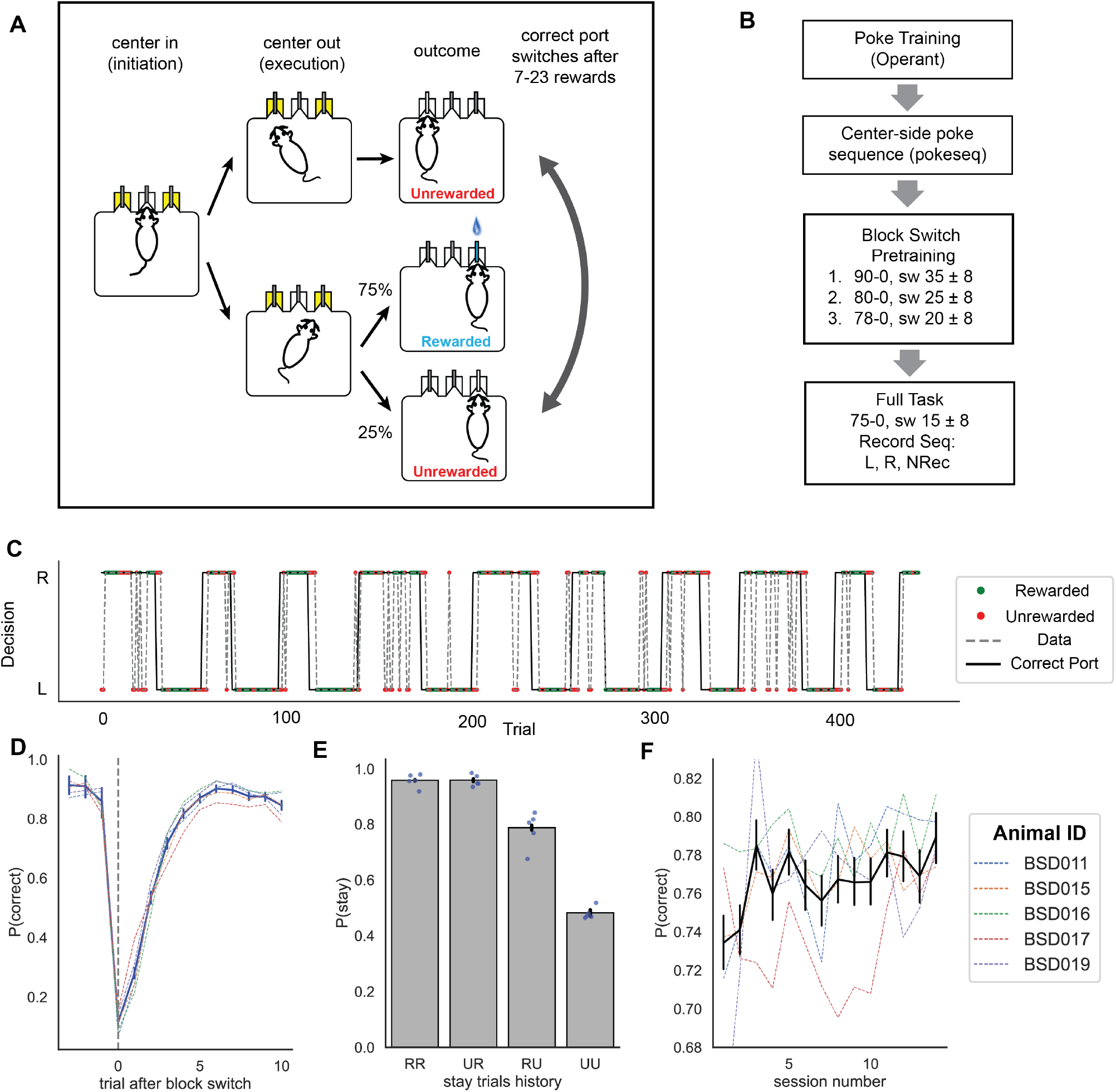
Mice adapt rapidly to block switches in a probabilistic reversal task. (A) Illustration of the two-armed bandit task, divided into initiation, execution, and outcome phases. In the illustrated trial, the right port is rewarded with 0.75 probability and the left port is unrewarded. After 7-23 rewarded trials, the correct port switches. (B) Training protocol. The recording phase took place in the “Full Task” phase. In the pretraining phases, the structure of the task was the same as the in the full task phase, except the reward contingencies and block lengths were different. Each contingency is labeled by numbers indicating the proportion of correct and incorrect choices that were rewarded. For example, “90-0” in the first pretraining phase indicates that 90% of correct choices were rewarded. The block length in each phase is indicated by its mean and range. For example, “sw 35 ± 8” in the first pretraining phase indicates that switches occurred after the animal earned between 27 and 43 rewards. During the 14 sessions of mice behavior data collection, we recorded dLight signals using a “left hemisphere (L), right hemisphere (R), no neural recording pure behavior (NRec)” sequence. (C) Raw behavioral trajectory taken from the first half of a sample session. Black line indicates correct reward port locations while dashed gray line indicates actual mouse behavior. Green dots and red dots mark rewarded and unrewarded trials, respectively. (D)Probability of making a correct choice (i.e., choosing the high probability port) as a function of the number of trials around a block switch. The vertical dashed line shows trials at which rewarded block changes. Each colored dashed line plot shows behavioral performance for individual animals. (E)Probability of staying (repeating the last choice) after experiencing different outcome histories in the same port. RR: two consecutive rewards; UR: unrewarded outcome followed by rewarded; RU: rewarded outcome followed by unrewarded; UU: two consecutive unrewarded outcomes. (F) Performance across 14 sessions. Dashed lines show individual animal trajectories. Error bars show 95% bootstrapped confidence intervals.

Another limitation of the existing literature is that many studies compare only a few model variants in isolation, without including a larger growing set of models. In this study, we included a broad set of complex RL models commonly used in cognitive modeling. In addition to comparing “pure” Bayesian and RL models, we can also examine hybrid models where Bayesian and RL processes are combined [9, 21, 22]. In these belief state RL hybrid models (BRL), Bayesian inference can be used to compute belief states, over which RL processes operate to learn policies appropriate for the current belief state.

Given these ambiguities in the past literature, our goal was to examine if Bayesian models or Bayesian-RL hybrid models could outperform sophisticated variants of RL models to explain behavior and outcome related dopamine signals in an instrumental task with a hidden state. We chose to focus on the nucleus accumbens (NAc) dopamine release because mesolimbic dopamine neurons have been widely associated with RPE predictions, one of their major projections destinations is the NAc, and the NAc is thought to play a role in the credit assignment [23–26]. To this end, we trained mice in a value based 2ABT task while recording fluctuations of dLight signals in the NAc using fiber photometry to measure dopamine transients[27]. These data allowed us to test the hypotheses about which computations mice use to solve the 2ABT at the behavioral level and to test if these computations are reflected by signals relayed to the NAc by midbrain dopamine neurons.

## Results

### Mice rapidly adapt to block switches in the 2ABT

To study the flexible updating of goal-directed behaviors under probabilistic conditions, we trained mice on a 2ABT task with block switches. Two weeks prior to training, mice were injected bilaterally with AAV dLight in the NAc core and implanted with an optic fiber and ferrule above the injection site.

In the 2ABT, mice encountered three ports equipped with infrared sensors. Animals were water restricted and trained to nose-poke into the central port to initiate a trial and then move to a left or a right port to obtain a water reward. Water rewards were available on either the left or right side, depending on the block. The correct choice was rewarded with water 75% of the time whereas no rewards were available at the other port. After they received a random number of rewards (uniformly sampled from 7 to 23 for the “Full Task” condition, Fig. 1A-B), the correct port switched without any discriminative cue. In order to achieve consistent water rewards across the whole session, mice needed to readily update their choice using outcome feedback.

As soon as the full task started (see Methods for details), we recorded unilaterally from the NAc in mice performing the 2ABT for a total of 14 daily sessions, alternating the hemisphere and allowing one non-recording session every third day (Fig. 1B). Just after pre-training, mice chose the high reward probability port 72.9% (95% CI: [0.669, 0.767]) of the time on average. Over the course of 14 training sessions, performance steadily increased (Fig. 1F, linear regression slope coefficient 0.0026, t statistic: 2.743, p=0.008, CI: [0.001, 0.005]). In the first 7 sessions, mice took 2.49 trials on average (95% CI: [2.39, 2.59]) to switch to the correct port after a block switch. In the 8-14th session this reduced to 2.36 trials on average (95% CI: [2.28, 2.45]). Mice switched and committed to the correct port for 3 or more trials (average 3.12 trials, 95% CI: [3.00, 3.26], Fig. S1) following the first 7 sessions. These data were comparable to mice performing the task without a fiber implant or cable [16].

To identify what task features the mice might consider before switching ports, we analyzed the stay/switch behaviors after two trials of outcome history. To simplify the comparison, we specifically chose the trials after the mice consecutively stayed in the same port for two trials or more. We encoded the outcome history at t-2, t-1 as RR, RU, UR, RU, (23607, 5719, 9428, 8520 trials each) with R representing rewarded trials and U representing unrewarded trials. For instance, a trial with “RU” stay history means that a mouse encountered a reward at trial t-2 and then got unrewarded at trial t-1, all at the same port. We found a significant reduction in the probability of repeating the previous choice after one unrewarded outcome (RR vs. RU, Fisher’s exact test: 6.47, p ≤ 1e-4, Cohen’s d: 0.6242, 95% CI: [0.6, 0.65]). Moreover, mice reduced their stay rate at the chosen port after two consecutive unrewarded outcomes compared to one unrewarded outcome (RU vs. UU, Fisher’s exact test: 3.98, p ≤ 1e-4, Cohen’s d: 0.6715, 95% CI: [0.64, 0.7]; Fig 1E).

### Cognitive model fitting confirms that mouse behavior data favors Bayesian models or complex RL models compared to simple RL models

To investigate possible computations that mice use to rapidly adapt their choices following block switches, we compared a purely Bayesian model (BIfp, see methods) adopted from [17], a BRL model (BRLfwr, see methods) inspired by [9], and several RL models (Fig. 2): a model with asymmetric learning rates for positive and negative RPEs (RL4p, dubbed the simple standard RL model), a model that additionally used counterfactual updates for unchosen option values (RLCF), the recursively formulated logistic regression (RFLR) model developed by [18], an RL model that simulates forgetting via decaying the Q value of unchosen options (RLFQ3p), and a dynamic learning RL model that adjusts negative learning rate adaptively based on outcome uncertainty (RL_meta) [20]. Though one can show its connection to Hidden Markov Models (HMM) under restricted conditions, the RFLR model was grouped with other RL models due to its mathematical equivalence to the forgetting Q-learning model [18, 28]. Modeling details can be found in the Methods and descriptions shown in Table 1. A summary of the model relationships is shown in Fig. 2A.

**Table 1:**
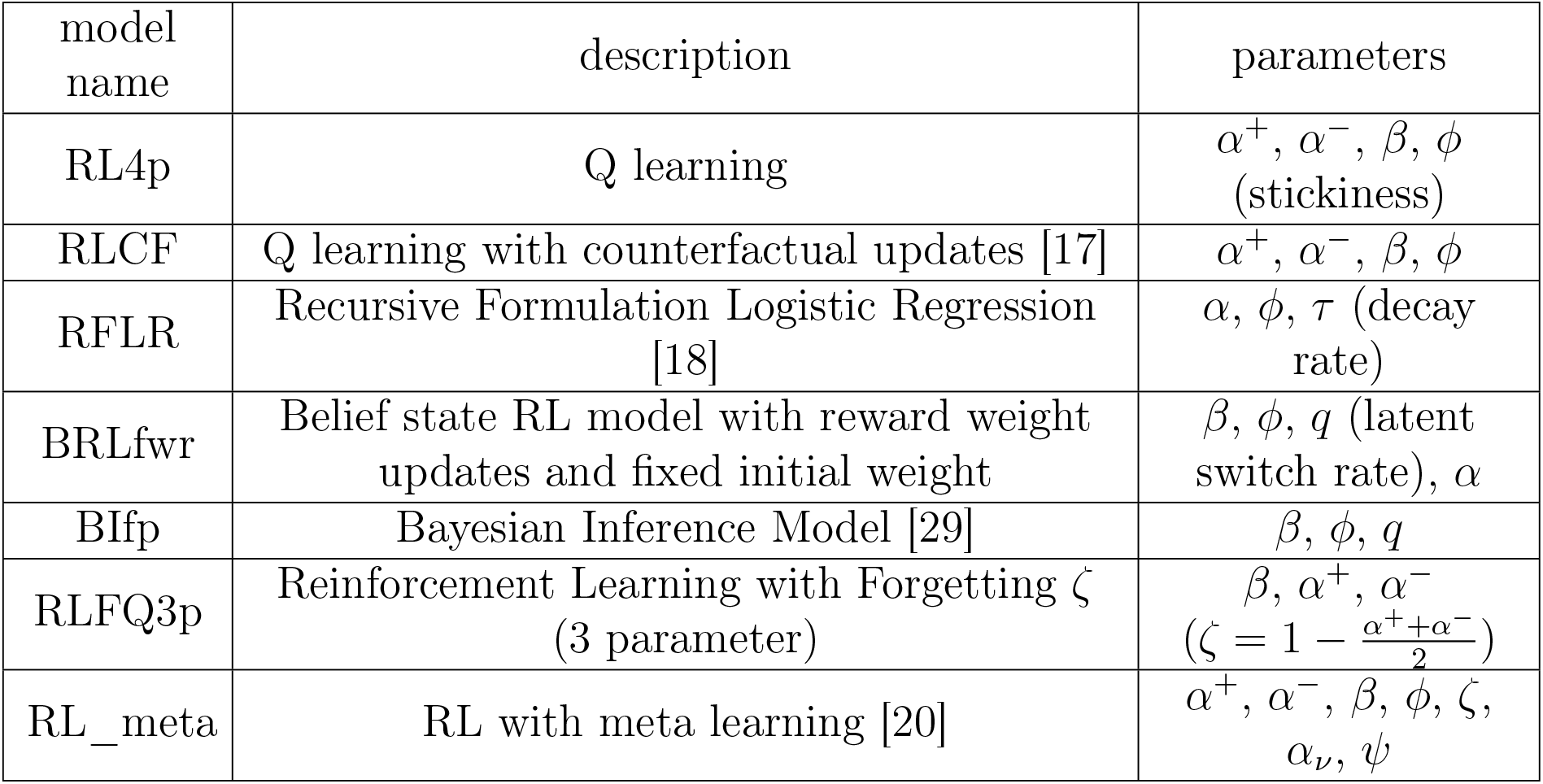
Overview of different cognitive models.

| model name | description | parameters |
| --- | --- | --- |
| RL4p | Q learning | $\alpha^+, \alpha^-, \beta, \phi$ (stickiness) |
| RLCF | Q learning with counterfactual updates [17] | $\alpha^+, \alpha^-, \beta, \phi$ |
| RFLR | Recursive Formulation Logistic Regression [18] | $\alpha, \phi, \tau$ (decay rate) |
| BRLfwr | Belief state RL model with reward weight updates and fixed initial weight | $\beta, \phi, q$ (latent switch rate), $\alpha$ |
| BIfp | Bayesian Inference Model [29] | $\beta, \phi, q$ |
| RLFQ3p | Reinforcement Learning with Forgetting $\zeta$ (3 parameter) | $\beta, \alpha^+, \alpha^-$<br>($\zeta = 1 - \frac{\alpha^+ + \alpha^-}{2}$) |
| RL_meta | RL with meta learning [20] | $\alpha^+, \alpha^-, \beta, \phi, \zeta, \alpha_\nu, \psi$ |

**Figure 2.**
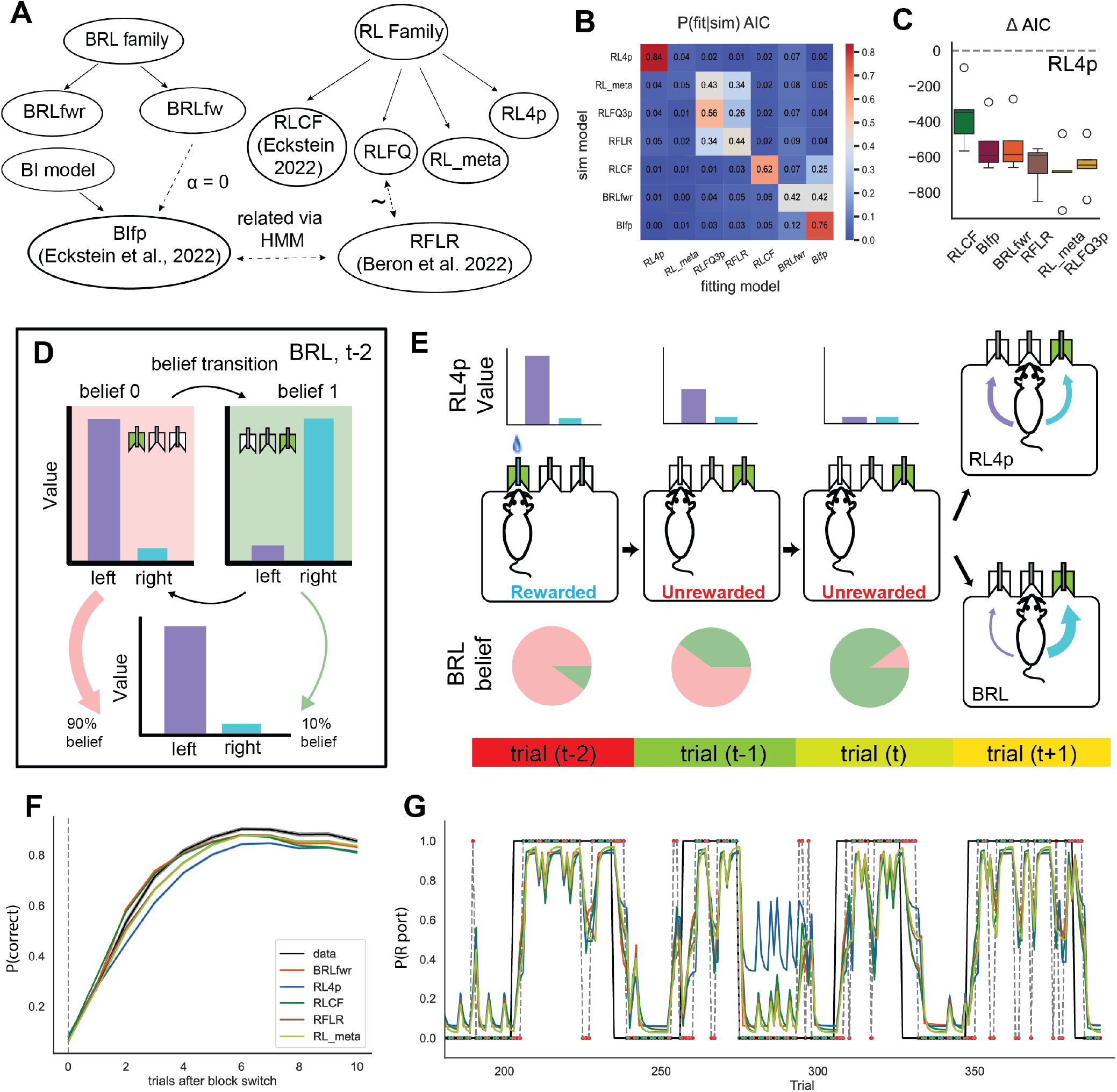
Bayesian and reinforcement learning models. (A) Relationships between cognitive models (see Methods for more details). (B) Confusion matrix outlines results for model identification analysis. Each entry i, j represents the percentage of time that the column j fitting model best explained data generated by row i simulating model. The row orders are sorted via dendrogram based on model similarity (see Methods). (C) Model comparison using relative AIC compared to RL4p: ΔAIC = AIC(model)-AIC(RL4p), with lower values indicating better fit. (D) Illustration of value computation for BRL model family, which updates beliefs via Bayes’ rule and then uses these beliefs to compute values. (E) Illustration using a four-trial sequence to show the differences between RL4p and BRL. Top: purple and cyan bars show the choice values conditioned on the belief state; Bottom: pie charts show the belief state for BRL; the animal’s policy is selected as a function of the value within their belief states. (F) Behavior of different models compared to mouse data (black line). Trial 0 is when the program has switched the rewarded side in a block switch. (G) Example behavioral trajectory (probability of choosing the rightward port) predicted by different models. Mouse data are marked by a dashed line and block structure is marked by a solid line. Rewarded trials are marked as green dots and unrewarded trials are marked as red dots. Error bars show 95% bootstrapped confidence intervals.

All models listed compute choice values (or implicit values via reward probabilities in the case of BIfp), map these to choice probabilities, and then update their values (and beliefs in the case of the Bayesian models) after receiving reward feedback. For models that do not explicitly utilize RPE for model updating, like BIfp, we can calculate “pseudo-RPEs” by taking the difference between the observed and expected reward. Critically, the BRL and RL models differ in how they perform value computation. The RL models update choice values stored in a look-up table, while the BRL models computed values as a sum of choice values in each belief states weighted by posterior belief probabilities. The updating process is schematized in Fig. 2D-E.

The RL4p model and its variants have been extensively used in previous studies (e.g., [30– 34]), which have provided evidence that asymmetric learning rates and choice perseveration (“stickiness”) are often helpful in capturing animal behaviors. RL4p serves as a baseline against which all more complex models should be compared. Despite its past empirical success, a critical limitation of the RL4p model is that it fails to capture the observation that animals appear to update values for unchosen options (counterfactual updating; [17, 35–37]). For example, in reversal learning tasks like the 2ABT, observing an unexpected reward omission dramatically increases the likelihood of a switch, accompanied by neural responses that anticipates the new reward contingencies [19, 38]. Counterfactual updating is usually formalized by updating values for unchosen actions in the opposite direction from the values of chosen actions. This qualitatively mimics the behavior of Bayesian models[29].

We fitted all models to the choice data of all 14 sessions for 5 mice (Fig. 1F) in the 2ABT using maximum likelihood estimation. To qualitatively analyze similarities among this rich set of cognitive models, we simulated behaviors using parameter ranges fit to the mouse behavioral data. Then we fit each of the 7 main model variants to each of the simulated behavioral data sets and tested for model identifiability based on the frequency with which the ground-truth model was chosen by the model selection criterion, the Akaike Information Criterion (AIC). This revealed relatively poor identifiability between the Bayesian models, as well as poor identifiability between the complex RL models (RL_meta, RLFQ3p, RFLR). These results indicate that the 2ABT cannot be used to discriminate within each family of model on the basis of behavioral data, but can discriminate across families. Furthermore, the 2ABT is adequate for rejecting the standard RL model (RL4p, Fig. 2B,C).

With RL4p as a baseline, we found that both BRLfwr and BIfp significantly improved the AIC measure compared to the RL4p model (BIfp: ΔAIC = −536.57 ± 59.48, BRLfwr: ΔAIC= −530.59±61.86). The RLCF also explained the mouse behavior better than the RL4p model (RLCF: ΔAIC = −361.92±70.52), with a higher relative AIC on average than the BIfp model (ΔAIC = 174.64±47.83) (Fig. 2C). Additionally, other complex RL models also outperformed the RL4p model (ΔAIC = −652.53±49.26, −683.58±61.31, −646.45±53.62 for RFLR, RL_meta, RLFQ3p respectively, Fig. 2C). When we evaluated the qualitative difference for model fitting across subjects, we noted that due to the limitation of sample size, the lowest attainable uncorrected p value against a one-sided alternative using nonparametric pairwise tests (like permutation test) is 0.03125, which can readily lose power when multiple test corrections are used. Furthermore, it is reasonable to assume that relative AIC values across subjects follow a symmetric and normal-adjacent distribution.Therefore, we used parametric t test and controlled family-wise error rate (FWER) via Holm’s method with Bonferroni adjustment. We found that both Bayesian models like BRLfwr (p corrected: 0.012) and BIfp (p corrected: 0.013), and complex RL models (p corrected for relative AIC for RFLR: 0.003, RLFQ3p: 0.004, for RL_meta: 0.005) provided better accounts of mouse behavior in the 2ABT than the standard RL model. There was not sufficient discriminative power, however, to identify whether Bayesian models or any member of the complex RL models was superior based on the simple quantitative account of all behavioral data as a whole (corrected p value for BRLfwr vs RL_meta: 0.20, BRLfwr vs RFLR: 0.370, BRLfwr vs RLFQ3p: 0.366, though BRLfwr is significantly lower than RLCF: p corrected: 0.015).

### Bayesian models and complex RL models qualitatively recover mouse behaviors around trials feature volatile outcome changes

To further understand the mechanism by which the Bayesian models or the complex RL models explained mouse behavior, we compared qualitative signatures of the BIfp, BRLfwr, and RL models with those of mouse data. One hallmark behavioral signature of mouse adaptation to block switches is improved choice accuracy as the number of trials after a block switch increases (Fig. 1D). Due to the model similarities found in the previous model identification analysis, (Fig. 2B,C), we focused on a restricted subset of model space: BRLfwr, RLCF, RL4p, RFLR and RL_meta. We omitted BIfp and RLFQ3p from the analysis due to their qualitative similarity to BRLfwr and RFLR respectively. Accordingly, we first compared the rate at which choice accuracy improved after block switches for simulated model behaviors against mouse data. Consistent with our hypothesis, we observed that BRLfwr and RLCF predicted faster adaptation to block switches compared to RL4p, resembling the adaptation rate of mouse switching behavior (Fig. 2F). When we focused on single-trial choice-outcome trajectories, we qualitatively observed that the Bayesian model and complex RL models updated their switch probability differently (or faster in most cases) from the RL4p model after unrewarded observations or near block switches (Fig. 2G). From this, one further hypothesis naturally arises: the Bayesian models and complex RL models outperform the RL4p model because of their ability to detect choice-outcome sequences that are suggestive of a block switch or volatile choice-outcome contingencies.

To test this hypothesis, it is imperative to obtain a more granular understanding of how different trial outcome histories drive a mouse to switch ports, and to determine if BRLfwr can successfully predict switch probabilities under different outcome histories, especially around block switches. Following [18], we categorized all trials based on their three past trial outcome histories, using capital A/B to denote rewarded outcomes and lower-case a/b for unrewarded outcomes. Due to the symmetrical task structure, we denoted the trial at t-3 as A/a regardless of its spatial location, and trials at t-2 or t-1 as A/a if the mouse chose the same port, or B/b if the mouse chose a different port, in reference to trial t-3 (Fig. 3A). For instance, if a mouse was unrewarded at port 1 at t-3, rewarded at port 2 at t-2, unrewarded again at port 2 at t-1, we would describe the trial outcome history as aBb. We then calculated the switch probability as the probability to choose a different port at trial t compared to trial t-1.

**Figure 3.**
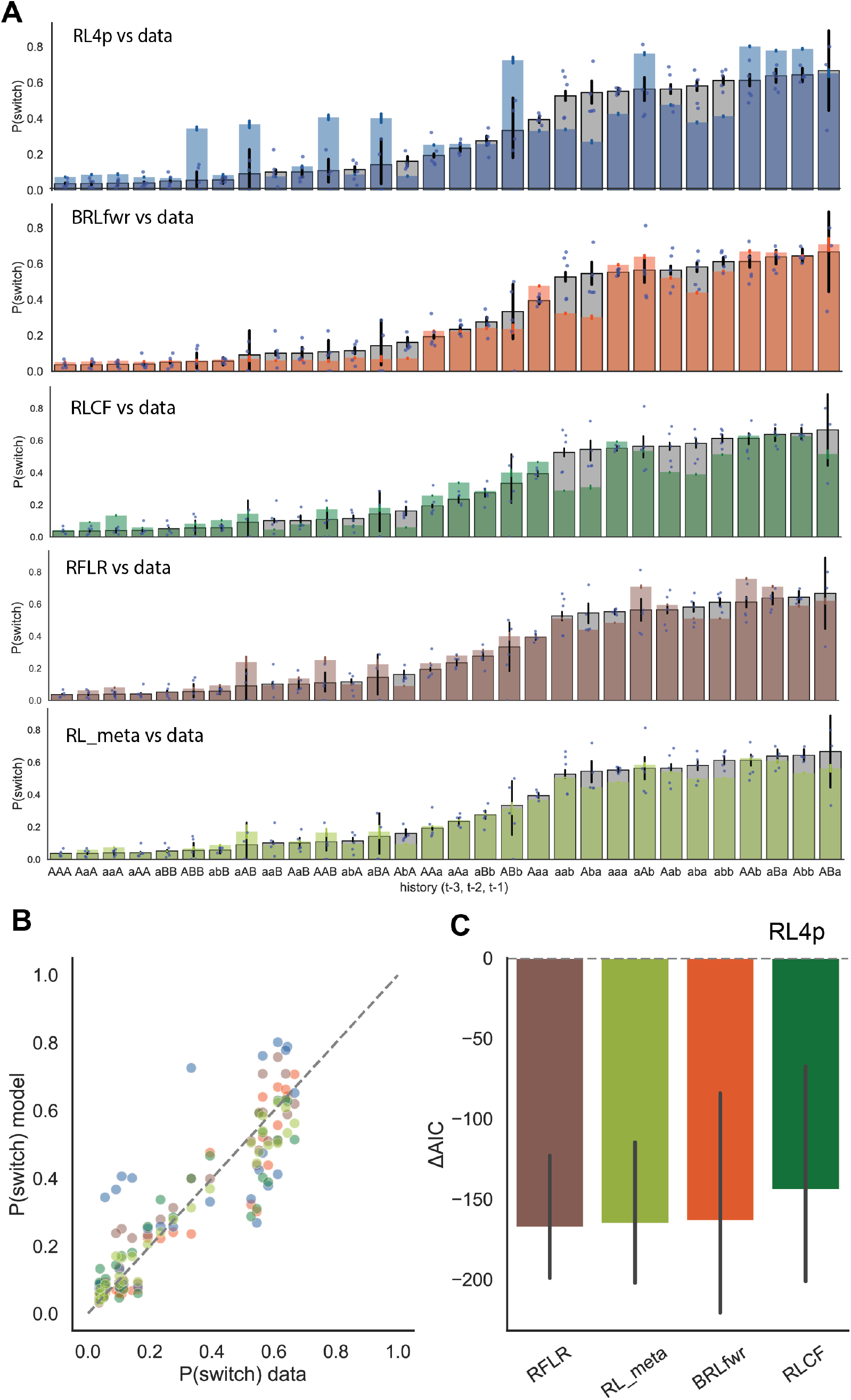
BRL and complex RL models outperform standard RL by better explaining mouse behaviors around block switches. (A) Switch probability by different trial outcome histories described by action-outcome pairs three trials back. Gray bars showed mouse average probability of switching for each outcome history, deep blue dots represent individual mice. From top to bottom: mouse data overlaid with BRLfwr, RL4p, RLCF, RFLR, RL_meta model predictions of switch rate, respectively. (B) Switch probability predicted by different models scales with probability of mice switching port selections in different outcome contexts described. Colors represent different models, sharing the same legend as C (orange: BRLfwr, wine red: BIfp, dark green: RLCF, brown: RFLR, blue: RL4p) (C) Relative AIC with respect to RL4p (dashed line at ΔAIC = 0) showing model fit to mouse data around block switch. Error bars show 95% bootstrapped confidence intervals.

We then compared the average mouse switch probability after each outcome history against the mean predictions of the three different cognitive models (Fig. 3B; sum of squared errors (SSE): BRLfwr: 0.1796, RL4p: 0.8270, RLCF: 0.2748, RFLR: 0.1537, RL_meta: 0.0787).

Notably, we observed that the RL4p model overestimated the switch probability for outcome history contexts where mice encountered rewards in both ports within the past 3 trials (e.g., ABb, AaB, etc., Fig. 3A). Since these outcome contexts occured only around block switches, mice usually stayed at the newly rewarded ports (mean switch rate: 0.2143). BRLfwr showed a similarly low switch rate after these contexts (0.1801), whereas RL4p model predicted a high average switch rate of 0.4328 (RLCF: 0.2248, RFLR: 0.2778).

For instance, after the outcome context of ABb, the RL4p model overestimated the probability of the mice switching back to port A (Fig. 3B), because it was only able to increment the action value for port B, instead of capturing the underlying reward distribution change. A mouse using a learning mechanism like RL4p would mistakenly think that selecting port A was still as valuable as it was prior to this outcome sequence, given that no reward omissions were recently experienced at that port, leading to regressive errors back to port A. Interestingly, the RL_meta model was able to explain adaptive behaviors with even higher accuracy than BRLfwr due to its adaptive power via learning rate tuning when reward context changes rapidly. We furthered our model comparison using relative AIC (ΔAIC) to the standard RL model (RL4p) for mice behaviors around block switch (defined as the first 5 trials after block switch), which showed similar ΔAIC results for BRLfwr and RL_meta (BRLfwr: -163.6763, RL_meta: -165.3471) (Fig. 3C).

Lastly, we noted that RL_meta, despite its great explanatory power, was also the most complex model with 7 parameters, with poor parameter identifiability (Fig. S2). In contrast, the more parsimonious Bayesian models exhibited good parameter recovery. To further investigate the differences between complex RL models and Bayesian models, we next turned to dopamine data analysis. We hypothesized that their mechanistic discrepancies would lead to qualitatively and quantitatively distinct RPE predictions, and henceforth distinct predictions of dopamine responses to action outcomes.

### Predictions from BRLfwr and BIfp capture nucleus accumbens core dLight dynamics better than RL models

We next investigated whether either Bayesian models or RL models provide qualitatively and quantitatively accurate predictions of dopamine release in the NAc triggered by rewards. Importantly, as noted above, BIfp does not inherently calculate an RPE term, since it uses Bayesian inference for model updates. Therefore, to allow the comparison between model predictions of BIfp and the dopamine signals, we calculated a pseudo-RPE for BIfp, the difference between observed and expected reward.

Our experimental mice were implanted with optical fibers bilaterally in NAc core to enable recording of dLight signals. Histology images were visually inspected after the experiment to verify the implant tip location and viral expression (Fig. 4A). We analyzed the dopamine responses in NAc by aligning dopamine signals to the “outcome” event, the time point when water rewards were either delivered to the mice or were omitted. After mice received a water reward, we observed a dLight dopamine signal (Z(DA)) increase on average (estimate: 1.6345, CI: [1.613, 1.656], p≤1e-5) in ports both contralateral and ipsilateral to the recording hemisphere, consistent with previous literature [7, 39, 40]. When reward was omitted at the peripheral port, we observed a reduction (OLS estimate: -1.5840, CI: [-1.604, -1.564], p≤1e5) in the dLight signal in NAc (Fig. 4B-C). These properties of NAc dopamine are broadly consistent with the RPE hypothesis: increases in response to surprising reward and decreases in response to surprising reward omission. Note that even when the mice have learned the underlying structure of the task, there is still irreducible uncertainty in reward delivery.

**Figure 4.**
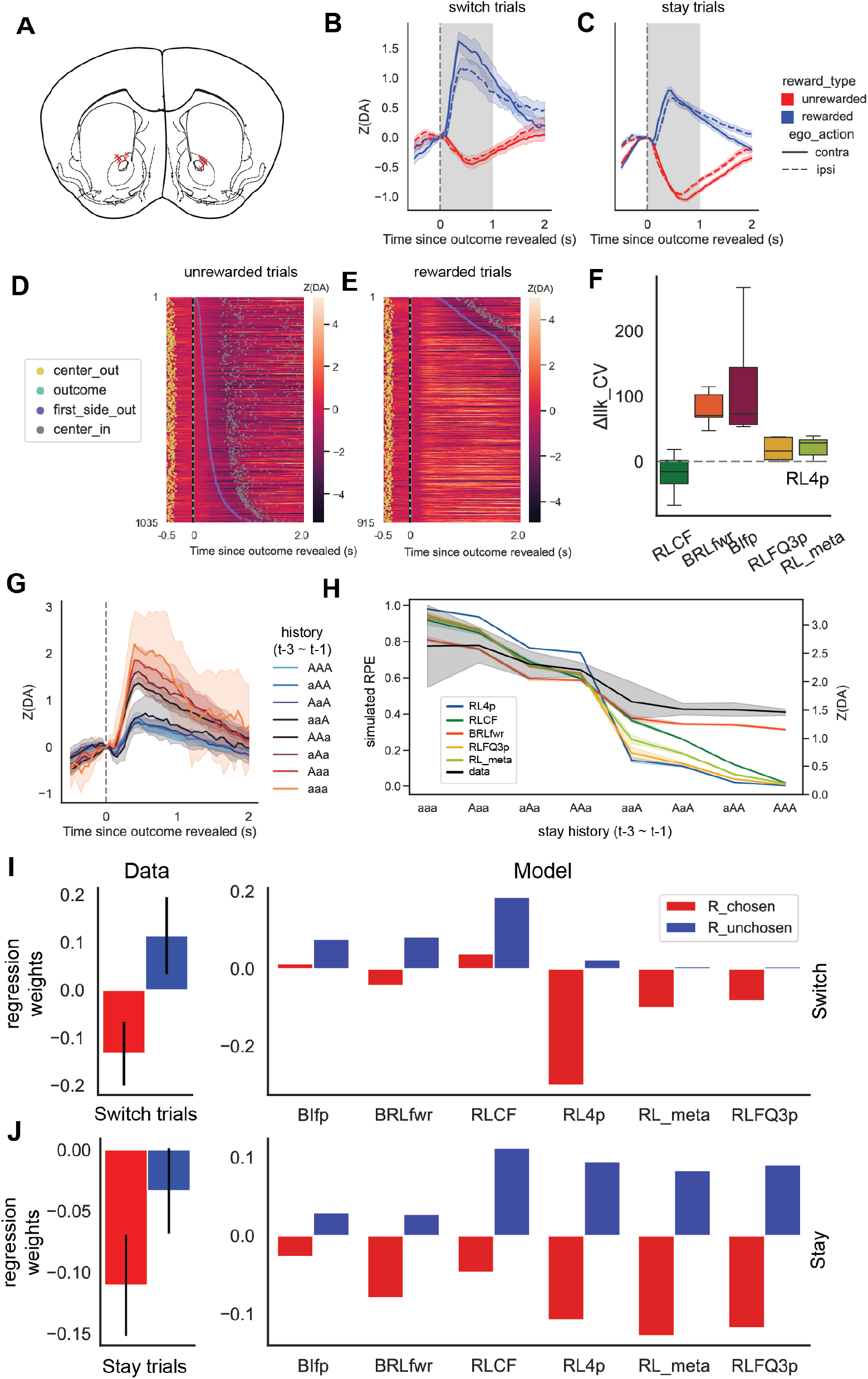
NAc dLight dopamine dynamics consistent with RPE predictions by models with Bayesian inference. (A) Implant fiber locations indicated on mouse brain atlas with red crosses. (B-C) Trial average of NAc dLight signals (z-scored, as described in Methods) aligned to outcome events. Shaded area indicates the one second where the peak or trough is taken for neural regression. (B) shows switch trials and (C) shows stay trials. Rewarded trials are in blue and unrewarded trials are in red. Trials where mice picked the port contralateral to the recording hemisphere are plotted with solid lines; trials in which mice picked the ipsilateral port are plotted with dashed lines. (D-E) Example session single trial dLight responses plotted in heatmaps, trials sorted by the time mice spent in the reward port (see Methods for further details). (D) shows a heatmap for unrewarded trials, and (E) shows rewarded trials. Increase in dLight signal is indicated by brighter shades of red and decreases from baseline are indicated by darker shades of black. Dots are used to mark “center out” (yellow), “outcome” (green), “first side out” (purple), “center in” (gray) events, respectively. (F) Result of neural regression using model RPE values to explain dopamine variability. Fit is measured as cross validated log-likelihood (llk_CV) relative to the RL4p model, with higher values indicating a better fit. Gray dashed line indicates the baseline of RL4p RPE fitted to dopamine measurements. (G) Dopamine response on rewarded trials binned by past history, sorted in increasing order of number and recency of rewards (note in all cases mice stayed with the same port ‘a/A’ for all three trials). (H) RPE predictions from different models plotted against dopamine peak values (in black). (I) Left: Relative change in dopamine as R_chosen (past rewards observed at the selected port) and R_unchosen (past rewards observed at the opposing port) change, calculated via LMER regression weights for dopamine observed in trials where the animals switched their port choices (animal switch trials). Right: Relative change in model RPE as R_chosen and R_unchosen change, calculated via regressions using model RPE predictions. (J) Similar to I, but for trials where the animal maintained their previous port selections (animal stay trials). Error bars show 95% bootstrapped confidence intervals.

To test the effect of animal choice switching, choice laterality (with respect to dopamine recording hemisphere), and reward or reward omissions on dopamine responses at outcome phase, we fitted a simple OLS model with all four factors. We observed a significant Z(DA) difference (OLS coefficient estimate for switch: 0.5030, CI: [0.454, 0.552], p≤1e-5) in trialaveraged dLight responses to the “outcome” event between switch trials and stay trials. Signals triggered by reward were larger on average on trials where mice switched to a new port, compared to when they stayed with their past choice. Negative signals observed after reward omissions in trials where mice stayed were larger when compared to when they had just switched (Fig. 4B-C). Additionally, our data are consistent with expectations that rewards are associated with increased dopamine release in NAc (estimate: -1.6345, CI: [1.613, 1.656], p≤1e-5) and unrewarded outcomes with a decrease (estimate: -1.5840, CI: [-1.604,-1.564], p≤1e-5). There was no significant effect of hemisphere (laterality, contra or ipsi relative to reward port) on NAc dopamine release (p =0.523).

When we visualized single trial dopamine traces around “outcome” events, we found that the temporal span of the dopamine response qualitatively aligned with the amount of time mice stayed at the peripheral reward port (Fig. 4E). Similarly, after unrewarded outcomes, we found that the duration of the decrease in dLight response qualitatively aligned with the time mice stayed in the peripheral reward port, reaching signal trough typically just after mice left the port. These results together suggest that when modeling dopamine responses, the time duration that mice stayed in the peripheral reward port after an “outcome” event (dubbed “port duration”) needs to be taken into consideration (Fig. 4D-E). For this reason, we included port duration as a covariate in our regression analyses.

We reasoned that if the internal value updates of the mice are computed by dopamine in the NAc and resemble that of BRLfwr or BIfp, then BRLfwr or BIfp RPE calculations should explain a larger amount of variance in the dopamine responses in NAc at the time of “outcome” events than RL models. To simplify the testing procedure, we obtained dopamine summary statistics by taking the peak of dopamine transients for rewarded trials and troughs for unrewarded trials (dubbed DA-PT) during the one second window after the outcome, marked in gray in Fig. 4B. To control for behavioral confounding variables, we included session number, port duration, egocentric action, movement time, and center port poke duration as covariates in addition to predicted RPE values from BIfp, BRLfwr, RL4p, and RLCF (note that we omitted RFLR model since it does not have a straightforward RPE formulation), each in a disjoint regression covariate set. Using maximum likelihood estimation with featurized covariates (see Methods) on the five different RPE covariate sets, we found that both BIfp and BRLfwr models yielded larger relative log-likelihood (Δllk CV) than the RL4p baseline, evaluated on cross validation sets (70-30) for each animal (Fig. 4F, Δllk CV: BRLfwr: 80.58, 95% CI: [60.84, 102.79], BIfp: 118.68, 95% CI: [65.71, 217.87]). Both Bayesian models also outperformed other complex RL models on this measure (Δllk CV: RLCF: -19.62, 95% CI: [-47.40, 3.64], RLFQ3p: 18.83, 95% CI: [4.97, 33.30], RL_meta: 22.22, 95% CI: [7.93,33.75]).

To illustrate the quantitative differences between model predictions, and to identify the aspects of the data that the BRL and BIfp models captured but the RL models did not, we investigated conditions where different models gave qualitatively different predictions. First, we looked at the changes in dLight responses to rewards as mice experienced a series of outcomes in consecutive stay trials from aaa to AAA (lowercase “a” indicates unrewarded and uppercase “A” indicates rewarded). As mice encountered more rewards in their recent history, dLight dopamine responses to reward gradually decreased (consistent with decreased RPE) but did not completely flatten to no response (Fig. 4G), partially explaining the quantitatively worse fit (fig. 4F). This pattern observed in our dLight data from the NAc core (‘data’ indicated by a black line) was not well captured by the RL models in later trials but was captured by the Bayesian models. To quantify, we trained a separate OLS model for each cognitive model to predict dopamine response using only simulated RPEs. We compared the model fitness across different cognitive models with cross validated loglikelihood (BRLfwr: -985.89, RL4p: -991.80, RLCF: -992.87, RLFQ3p: -994.74, RL_meta: -995.02) (Fig. 4H). Indeed, BRLfwr RPE predictions achieved the highest model fitness to dopamine data. The RL models updated action values on each trial as mice encountered consecutive rewards, estimating a recent average of choice outcomes. This feature may make RL models overly sensitive to recent outcome histories and liable to overestimating the action values, particularly around block switches. In our dataset, the best fitting RL4p model had an average *α*^+^ of 0.85 (95% CI: [0.77, 0.94]), and average *α*^−^ of 0.73 (95% CI: [0.71, 0.75]). The large *α* (learning rate) values likely enabled RL4p to maintain the flexibility to adapt to block switches, but they also posed issues regarding biological feasibility. RL_meta was able to mitigate this issue with a dynamic learning rate that adapts to outcome uncertainty. However, due to its iterative nature, as the amount of reward in the recent outcome history increased, it steadily decreased the RPE towards zero, which did not match what we observed in the dopamine signal.

Both BIfp and BRLfwr were able to explain the adaptation of dopamine transients after increasing consecutive rewards better than the RL models. These models accounted for possible block switches and maintained values of both a high reward context and a low reward context that are not associated with any given port (Fig. 4H). Instead, these values were assigned to specific ports in proportion to the model’s belief state on any given trial. We speculate that this information about task structure provided robustness in response to probabilistic reward omissions after selecting the more rewarding port. This also allowed the best fitting BRLfwr and BIfp model to predict a small but non-negative RPE response on rewarded stay trials (Fig. 4H), suggesting a degree of uncertainty as to whether the block had switched and capturing a pattern observed in the NAc dLight signals in later consecutive trials (right side of Fig. 4H). Furthermore, we observed that among trials after an immediate past trial reward, the dopamine response was highly consistent, independent of more distant outcome histories. Only BRLfwr captured this phenomenon qualitatively, in striking contrast to other complex RL models.

The Bayesian models were able to modulate expectations about both choices using inference based on one single choice outcome. This leads to the prediction that at the outcome phase of each choice, an increasing number of rewards gathered at the opposing port in past trials will reduce the RPE signal reflected in dopamine, especially trials where animals switched to a different port from the previous trial (animal switch trial). To investigate if this prediction was upheld in the dopamine data, we fitted both the mouse data and the model simulated data with Linear mixed effects models (LMER, see Methods). Specifically, we modeled the dopamine values, or the predicted RPEs for simulated data, as a linear sum of Reward, R_chosen (past rewards observed at the selected port) and R_unchosen (past rewards observed at the opposing port), conditioned on animal switch trials or animal stay (trials where the animals maintained their prior choice) trials. On animal switch trials, both BIfp, BRLfwr, and RLCF predicted a positive effect of R_unchosen, but other RL models predicted little to no effect due to the forgetting effect or asymmetrical value updates. LMER fitting revealed a significantly negative effect of R_chosen on dopamine at outcome phase during animal switch trials (slope estimate: -0.128, 95% CI [-0.195, -0.062], p=0.02), and a significantly positive effect of R_unchosen on dopamine during animal switch trials (slope estimate: 0.126, 95% CI [0.035, 0.217], p=0.03, Fig. 4I). On animal stay trials, the result was more complicated. All models predicted a negative relationship between R_chosen and RPE signal, matching the dopamine observation (slope estimate: -0.09, 95% CI: [-0.11, - 0.07], p=0.01). However, low sample size of high R_unchosen value trials a noisy negative estimate of effect of R_unchosen, which none of the models generated consistent predictions (Fig. 4J). This elevated level estimation errors for R_unchosen makes it a bad target for model arbitration, while the other three measures may be a more credible source of evidence. To sum up, the relationship between past reward history and dopamine values qualitatively match the prediction of the BRLfwr model for switch trials and the predictions of BIfp or RLCF model for stay trials. Taken together with the functional similarity of BIfp and BRLfwr (Fig. 2B), we conclude that the dopamine data matched the RPE predictions of Bayesian models quantitatively and qualitatively, but we could not further discriminate between BIfp or BRLfwr.

## Discussion

Our results showed that mice were capable of rapidly adapting to block switches in an instrumental task with probabilistic and volatile contingencies, without sacrificing the robustness of their choice policy when encountering stochastic unrewarded outcomes after correct choices. Both Bayesian models (the pure Bayesian inference model and the belief state RL model) were able to capture this balance and outperformed RL models in explaining mouse behavior (Fig. 2C, Fig. 3C) and dopamine release events in the NAc core following choice outcomes (Fig. 4G-J). Critically, both Bayesian models had explicit belief state representations of different block identities (Fig. 2DE), which provided a mechanism for encoding the rapidly changing reward contingencies in the 2ABT. This mechanism successfully explained the ability to efficiently switch after consecutive unrewarded outcomes while also disregarding occasional reward omissions within a block. The RL component in the BRL models allowed the model to use RPEs to update its mapping from choices to values conditioned on belief states. However, we did not find decisive evidence that BRL models explained the mouse data better than pure Bayesian models. The principal conceptual advantage of BRL models is that they make direct use of RPE signals, whereas pure Bayesian models do not. This highlights the importance of neural data in adjudicating between these models.

Building on the behavioral modeling, we showed that the Bayesian models do a good job predicting NAc dopamine activity at the time of outcomes (reward or reward omission). In particular, RPEs generated by the BRL model and pseudo-RPEs derived from the BI models qualitatively and quantitatively match the history-dependent pattern of dopamine activity (Fig. 4F-J). This finding supports a growing literature showing that dopamine signaling of RPEs depends on a belief state (or some approximation of a belief state; see [15]) in partially observable tasks [9–13]. Furthermore, our results show that in the context of a probabilistic instrumental task, VTA dopamine neurons compute RPE-like signals not just based on observable input, but also internally generated state information. This confirms an expanding literature showing that “model-free” RL computations in the midbrain and striatum take in higher-level inputs such as beliefs about latent state [9, 19, 22, 41], as well as information about reward outcomes and reward expectations [42–45].

The BRL model is closely related to structure learning models that learn multiple contextdependent policies. Such models often assume some form of Bayesian inference at the more abstract level, with RL supporting learning of specific policies [21, 46, 47]. Theoretical work, supported by experimental findings, has also shown that dynamic adaptation at the more abstract level of belief states/latent contexts/rules may also be supported by RL-like computations [29, 48, 49]. The current work cannot dissociate these possibilities.

There are several limitations to our work that can be addressed in future investigations. Here we adopted a 75-0 reward probability design from the legacy of prior lab literature. In this design, in any given block, an inferior port always yields no reward. Consequently, mice rarely switch to a new port and get rewarded after observing rewards in the previously chosen port. Previous work has noted that inference-based cognitive models are capable of decreasing the expectations about alternative options after observing a reward in selected option [19, 38, 41]. Therefore, mice using inference would exhibit a higher RPE following a reward in a new port after also observing a reward in the old port, a reward-switch-reward sequence. Yet in our dataset, there are only 94 trials initiating such sequences out of 30,707 trials in total, across multiple animal sessions, giving us limited power to address this prediction.

In our task design, mice can initiate new trials at their discretion after the outcome, with no instructed delays. Especially after unrewarded outcomes, mice leave the peripheral port shortly after, approaching the center port for a new trial. As a result, in trials when mice leave the peripheral port too soon, dopamine responses coincided temporally with the dopamine ramp associated with approaching center ports [50, 51]. To mitigate this, we only analyzed trials with specific port durations (see Methods), which limited our statistical power. More generally, the scaling effect of port duration on dopamine results could be another piece of evidence for functional diversity of dopamine neurons [52–57]. Moreover, our work does not directly address the possibility of other alternative computational hypotheses about dopamine, like directly setting the adaptive learning rate [58] or signaling retrospective inferences about causal targets [59].

Another limitation of our study is that the RPE that correlates with dopamine signaling only plays a minimal role in predicting behavior, as evidenced by the fact that BRL models did not outperform pure Bayesian models with no RL component. We believe that better discrimination between these models can come from theory-guided experimental design, possibly using a more complex task that places a greater demand on learning policies within each belief state [60, 61].

## Conclusion

In natural environments, animals and humans often experience ambiguous outcome feedback about the hidden reward structure of the environment. We emulated this in a two-armed bandit task with switching reward blocks, and showed that mice are capable of adapting rapidly to changes in hidden reward structures after observing specific outcome feedback sequences. Computational model fitting to behavioral data suggested that models performing Bayesian updating of beliefs better explained mouse behavioral data than standard RL models. These Bayesian models also quantitatively and qualitatively matched the dopamine release in the nucleus accumbens core better than the non-Bayesian alternatives. Together, we conclude that probabilistic belief updates are critical to behavioral adaptation and RPE signaling in the mesolimbic dopaminergic system during instrumental learning in a partially observable environment.

## Author Contributions

Conceptualization, M.K.E., A.G.E.C., S.J.G., C.D.H., A.J.Q., K.M., A.F.M., L.W. ; Methodology, S.J.G., A.J.Q., C.D.H., A.G.E.C, M.K.E, L.T., L.W., W.C.L. ; Software, A.J.Q., S.J.G. ; Validation, K.M., A.F.M., C.D.H., L.T. ; Formal Analysis, A.J.Q., S.J.G. ; Investigation, A.J.Q., C.D.H., E.M.T., L.T. ; Resources, L.T., L.W. ; Data Curation, A.J.Q., E.M.T., L.T., J.B.C. ; Writing - Original Draft, A.J.Q., E.M.T., L.W., S.J.G., C.D.H. ; Writing – Review & Editing, A.G.E.C., S.J.G., L.W., C.D.H., M.K.E., A.J.Q., L.T., W.C.L. ; Visualization, A.J.Q., S.J.G., A.G.E.C., L.W., C.D.H., K.M., A.F.M., M.K.E., W.C.L. ; Supervision, L.W., S.J.G., A.G.E.C. ; Project Administration, A.J.Q., C.D.H., L.W., and S.J.G. ; Funding Acquisition, L.W., S.J.G.

## Acknowledgements

We are grateful to Scott Linderman, Bernardo Sabatini, Celia Beron, Jing Jing Li, Thomas W. Elston, Yilan Liang, Hongli Wang, Gaia Molinaro, Jaclyn Essig, and Eric J. Hu for discussion and to Chris Machle, Lexi Z. Zhou and Anthony T. Dunn for discussion and assistance with data collection. We thank Prof. L. Tian for the provision of dLight1.2 AAV. This work was supported by the National Institutes of Health (grant number U19NS113201 to L.W. and S.J.G.) and the Air Force Office of Scientific Research (grant number FA9550-20-1-0413 to S.J.G.).

## STAR Methods

### RESOURCE AVAILABILITY

#### Lead contact

Further information and requests should be directed to and will be fulfilled by the lead contact, Albert J. Qü.

#### Materials availability

No newly generated materials associated with the paper.

#### Data and code availability

- All dLight fiber photometry recordings and mouse behavior data generated in this paper will be deposited in Open Science Framework in NWB format upon review, and will be publicly available as of the date of publication. Accession numbers are listed in the key resources table.
- All original code, including notebook used to generate figures, will be available publicly via github, further formulated upon review.
- Any additional information required to reanalyze the data reported in this paper is available from the lead contact upon request.

## EXPERIMENTAL MODEL AND STUDY PARTICIPANT DETAILS

### Animal protocol

Mice (all C57 Bl/6 male, bred in-house aged 104-167 days) were housed on a 12 h reversed light-dark cycle (lights on at 22:00) with nesting material. Prior to surgery they were group housed with access to food and water ad libitum. We conducted all animal procedures, which follow the principles outlined by the NIH Guide for the Care and Use of Laboratory Animals, according to the protocols approved by the University of California, Berkeley Institutional Animal Care and Use Committee (IACUC) and Office of Laboratory Animal Care (OLAC).

## METHOD DETAILS

### Surgery protocol

Mice were anesthetized with isoflurane gas for stereotaxic surgery. Meloxicam was given on the day of surgery and daily for 48h after. Coordinates for the Nucleus Accumbens Core were bregma coordinate: 1.20mm anterior, ± 1.2mm medial-lateral, -4.1mm ventral. AAV-CAG-dLight1.3b or AAV9-syn-dLight1.2b were injected using a Nanojet II (Drummond scientific). Neurophotometrics NA 0.37 or 0.48 400*µ*m optic fibers were placed approximately 100*µ*m above the injection site. Dental cement was used to secure the implant to the skull. During a recovery period of 7-14 days, mice were singly-housed and fed ad libitum.

### Histology and Imaging

Following behavior, animals were transcardially perfused with 4% paraformaldehyde (PFA) in an 0.1M phosphate buffer (PB) solution (pH = 7.4). Brains were collected and fixed overnight, followed by a transfer to 0.1M PB. To visualize striatal photometry fibers, perfused brains were sliced coronally at 50*µ*m using a vibratome (VT1000S Leica Biosystems; Buffalo Grove, IL). Immunohistochemistry was performed to amplify the GFP signal (1:1000 chicken anti-GFP, Aves Labs, Inc.; GFP-1020 followed by 1:1000 goat anti-chicken AlexaFluor 488, Invitrogen by Thermo Fisher Scientific; A11039). Slides were mounted on slides with Fluoromount-G (Southern Biotech). Slices anterior and posterior to the fiber tract were imaged at 10X on an AxioScan Z.1 fluorescent microscope (CRL Molecular Imaging Center, UC Berkeley) to confirm targeting.

### Probabilistic Switching 2ABT Task

We trained 5 male mice on a 2ABT behavioral task in which the location of the water reward was periodically switched between the two potential rewarded ports at random intervals. Trials were initiated by a nose poke in the center port and concluded after the mouse subsequently chose one of two reward ports, located on either side of the center port. Nose pokes were detected by a infrared photodiode. Once the initiating poke was sensed, lights that encompass the reward ports were triggered to cue the animals to the viability of a potential water reward. The mouse receives a 2 μl water reward or no reward depending on their correct or incorrect, respectively, port choice for each trial.

Only one peripheral port was rewarded at a time. The setting for the number and frequency of water rewards was altered during each phase of the task. Teaching animals the task, we started them in the “operant” phase, where water rewards were offered in the peripheral ports, and then moved to the “pokeseq” phase, where mice learned to poke the middle port before going to retrieve a peripheral port water reward. Successful learning during this phase was measured as the mouse getting more than 700 rewards in less than five hours. Some food pellets were placed into the arena while the mice were learning the task to motivate them. We omitted mice from the experiment if they didn’t perform successfully on “pokeseq” after five days.

Once the mice learned to poke the center port to initiate the peripheral port water rewards, we no longer put food into the arenas and advance the mice onto the “learning switch” phase. During the three subphases of the learning switch, we teach the mice that only one port is rewarded at a frequency of 90%, 80%, and 78% for each subphase respectively. Additionally, we taught the mice that the rewarded port switches after obtaining a random number of cumulative rewards set within the ranges of 27-43, 17-33, and 12-28 for each subphase respectively. Water rewards became less predictable and more variable with each subsequent phase.

Successful learning for the three learning switch phases were set at a score higher than 69.5% on both the left and right port within four, three, and two hours for each subphase respectively. If the mice seemed unmotivated, we placed some food into the arena. Once they completed the task with food, we removed the food and had them reach criterion for their appropriate subphase.

Fibers were first introduced to the mice after they complete the last learning switch subphase and are utilized for neural recording only after the mice successfully reach the learning switch criteria once again. During the “full task”, used for fiber photometry recording, mice received a water reward with a 75% chance if they picked the correct port, but received nothing if they picked the wrong port for each trial. Once mice obtained a random number of cumulative rewards, sampled from 7 to 23 uniformly, from one port, the reward was switched to the other port. This structure minimized the possibility for the mice to predict the timing of the switch.

When participating in this task, the mice were water restricted but maintained ad libitum access to food. They were supplemented with additional water if they earned less than 500 μl water from the task. Mice were weighed daily, before and after the task, and were given additional water if needed, at least 30 minutes after training sessions, to ensure that they maintained around 85% of their original weight.

### Fiber Photometry (FP)

We recorded from all 5 mice with fiber implants in nucleus accumbens core (NAc) using Neurophotometrics FP3001 system at 20Hz. dLight signals are measured in 470nm channel, while isosbestic control signal is measured in 415nm channel for artifact removal. During the learning switch phase, we start plugging in fiber implant without recording to get mice accustomed to moving and behaving with fibers. Starting from the first “full task” session, we alternate between a left hemisphere recording, right hemisphere recording, and no recording schedule. All FP recordings are synchronized to behavioral and video data via custom TTL systems and bonsai programs and saved for further processing. Recorded signals are preprocessed using custom python program to control for motion artifacts using 415nm reference channels [62]. Specifically, we first subtract the common DC shift by approximating a smooth trend across both channels. Then we perform a robust linear regression from reference channel to signal channel, and subtract the fitted 415nm artifact signal from 470nm channel recordings and obtain the processed dLight signals. All dLight signals are subsequently z-scored to account for across-session extraneous variations.

### Task Behavior Analysis

All final behavioral analysis is conducted with custom implemented python packages. Behavioral data are first collected using a custom implemented 2ABT control module in matlab, and then preprocessed into behavioral data frames with each row representing a separate trial with various columns containing information regarding task meta data, mice choice data, and task relevant variables. For trials that mice failed to initiate and make a choice, we included the data but noted the relevant variables as NaNs/null. For most behavioral analysis, outcome histories as well as covariates with null entries are dropped. During cognitive modeling, for null trials, we simplied maintained the model latent and skipped the model update steps. All model plotting are generated via seaborn packages, and test statistics and various measures are generated using statsmodels, pingouin and sklearn.

We divided the tasks into initiation phase, execution phase, outcome phase and inter trial interval (ITI) phase. The initiation phase includes both a “center in” and “center out” event when mice poke their nose into and out of the center port to initiate a trial. To execute a choice, mice leave the center port and then enter one of two peripheral ports to make a selection. When the outcome of their choice is revealed to the mice, they linger at the port for water reward or time-out until they leave the port during the “side out event”.

We defined the following critical events:

- Center in (CI): when mice first poke into the center port to start a new trial.
- Center out (CO): when mice leave the center port to select ports.
- Side in (SI): when mice poke peripheral port.
- Outcome (O): when outcome is revealed, typically immediately after side in (latency on the order of system latency).
- Zeroth side out (SO0): first time when mice move the nose poke out of the port trigger IR beam break.
- First side out (SO1): last time of a consecutive set of beam breaks at the same side peripheral port. Zeroth side out and first side out can often be the same when the animal leaves the port only after drinking all the water.
- Last side out (SOf): last side out beam breaks before next center port poke to initiate a new trial.

We defined the following critical timing measures:

- Movement time (MVM)T: CI - SOf (t-1)
- Inter-trial interval (ITI): CI - SO1(t-1)
- Center duration (center_dur): CO-CI
- Port duration (port_dur): SO1-O
- Side out bout latency (SO_lat): SO1-SO0, typically close to 0, critically this is used to control for atypical trials with excessive beam breaks.

### Computational Modeling

We formalize the decision problem facing animals as follows. At time *t*, the animal makes a choice *c*_*t*_ ∈ {−1, 1} and then observes a reward *r*_*t*_ ∈{0, 1}. By convention, we take *c*_*t*_ = −1 to denote the left port, and *c*_*t*_ = 1 to denote the right port. The reward probability depends on the chosen action and the hidden state *z*_*t*_ *∈* {−1, 1}:

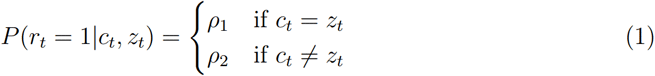

where *ρ*_1_ is the probability of reward for a correct choice, which equals 0.75 during the full task, and *ρ*_2_ is the probability of reward for an incorrect choice, which is technically 0 but which we set to 0.0001 to allow for model misspecification. The hidden state changes on each trial with probability *q* = *P* (*z*_*t*_*/*= *z*_*t*−1_).

We assume a common functional form for the choice policy across models:

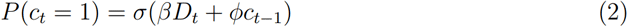

where *σ*(*x*) = 1*/*(1 + *e*^−*x*^), *D*_*t*_ = *Q*_*t*_(1)−*Q*_*t*_(−1) is the value difference, *β ≥* 0 is an inverse temperature parameter, *ϕ≥*0 is a stickiness (choice perseveration) parameter, and *Q*_*t*_(*c*) is the value assigned to choice *c* at time *t*. The stickiness component captures the tendency to repeat choices independent of the reward history.

The models make different assumptions about how the values are updated based on experience:

- **RL4p** is a standard RL model that updates Q-values according to a delta rule, with separate learning rates for positive (*α*^+^) and negative (*α*^−^) reward prediction error (RPE):

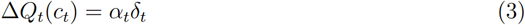

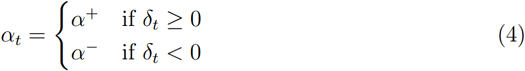

where δ_*t*_ = *r*_*t*_ − *Q*_*t*_(*c*_*t*_) is the RPE.

- **RLCF** is an elaboration of the RL4p model that uses “counterfactual” updates [17]. Whereas RL4p only updates the chosen action value, RLCF additionally updates the unchosen action value in the opposite direction:

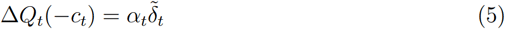

where −*c*_*t*_ is the unchosen action and 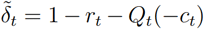 is the counterfactual RPE.

- **BRL** (belief state reinforcement learning) is an RL algorithm that computes the values as a linear function of the belief state *b*_*t*_(*z*) = *P* (*z*_*t*_ = *z*| *c*_1:*t*−1_, *r*_1:*t*−1_), the posterior probability over the hidden state given the choice and reward history (the belief state update will be described further below):

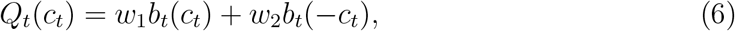

where *w*_1_ and *w*_2_ are learnable weights. Intuitively, the action value *Q*_*t*_ for a given action *c*_*t*_ is composed of two terms: the first term captures the reward weighted by probability when the animal is correct (*z*_*t*_ = *c*_*t*_), and the second term captures the reward weighted by probability when the animal is incorrect (*z*_*t*_ = −*c*_*t*_). If the animal has perfectly learned the task, the weights should be dictated by the ground truth reward probabilities, *w*_1_ = *ρ*_1_ and *w*_2_ = *ρ*_2_. We formalize a model of learning (**BRLfw**) based on gradient descent, which allows the animal to approximate task rewards without knowing ground truth:

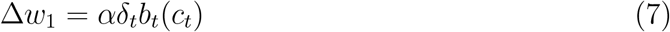

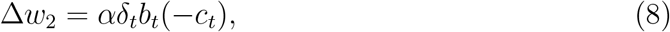

where *α* is a learning rate. We considered two versions of the model. The weights were initialized to *w*_1_ = *ρ*_1_ and *w*_2_ = *ρ*_2_ (note that choosing other initial conditions had relatively little effect on model fits). Note that the weights can be interpreted as representing the Q-values of correct actions conditioned on the belief, with their update rule a confidence-weighted RL update (see e.g., [63]), hence the name BRL. We also considered a restricted version of the model (**BRLfwr**) where we fix *w*_2_ = *ρ*_2_, which we found to perform just as well as the unrestricted version, so we focused on the restricted version in the main results. Belief state updating followed Bayes’ rule, with 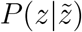 denoting the transition probability of the hidden state from 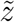to *z*:

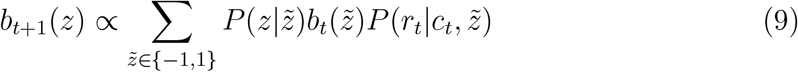

We assume that the belief state is initialized to *b*_1_(*z*) = 0.5. Also we parameterize 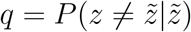 as the transition probability parameter of the model.

- **BIfp** (Bayesian inference with fixed parameters) makes use of the same belief state as the BRL models, but where there is no updating of the weights. Essentially, *D*_*t*_ = 2*b*(1) −1. This is equivalent to *w*_1_ = *ρ*_1_ and *w*_2_ = *ρ*_2_ for BRL. Interestingly, the best fitted parameter for *α* in the unrestricted BRL version (BRLfw) above was zero for most subjects, establishing an equivalence, under this 2ABT task, between BIfp and and BRLfw.
- **RFLR** (recursively formulated logistic regression; [18]) is a form of RL that is mathematically equivalent to a form of logistic regression. RFLR updates the value difference variable according to:

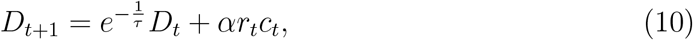

where *τ* is a timescale parameter governing the exponential decay rate of the values. Notably, we removed the non-negativity constraint on *ϕ* and fixed *β* = 1 in equation 2 for RFLR specifically, as it is consistent with the definition in [18] and yields slightly better performance.

- **RLFQ3p** (reinforcement learning model with forgetting, totaling 3 parameters) is a popular RL-based framework to model the forgetting, or Q value decay, of the nonchosen options. The formulation is similar to standard reinforcement learning model. However, Q value of the unchosen option decays as *Q*_*t*_(−*c*_*t*_) = ζ*Q*_*t*−1_(−*c*_*t*_). For model simplicity, we take 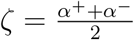. Interestingly, the forgetting parameter ζ captures stickiness effect of past chosen options, as one could show their mathematical equivalence. Consequently, the RLFQ3p model we used here has low model complexity with only 3 parameters *β, α*^+^, *α*^−^.
- **RL_meta** (reinforcement learning model with “meta learning” [20] adopts a similar general framework as RL model with forgetting (ζ is the forgetting parameter), but adapts the learning rate parameter by unexpected uncertainty *ν* (the rate of this adap-tion is controlled via *ψ*). In other words, *α*^−^ varies, subjected to non-negativity, as a function of how surprising recent outcomes were:

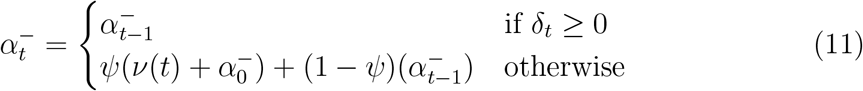

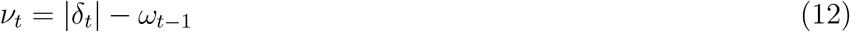

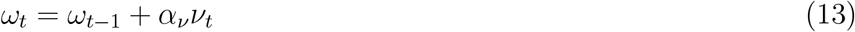

Moreover, expected uncertainty variable *ω*, initialized from 0, further mediates the update of Q values as following, on top of learning rate adaptations:

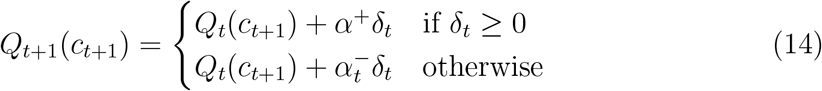

### Neural Data Analysis

All analysis was carried out with a custom Python module for aligning, visualization, and data modeling. After baseline correction, we visually inspected both 415nm channel recording and 470nm channel recording, as well as the AUC-ROC scores between the rescaled baseline values against signal channel recordings. We picked sessions that meets the following conditions: AUC-ROC higher than 0.9, longer tail in distributions in 470nm recording compared to rescaled 415nm recordings suggesting clear fluorescence increases, and no sharp discontinuity in both channels’ recordings. We use baseline-corrected and z-scored dLight signals for in-depth analysis, called Z(DA). For neural data alignment, we interpolated Z(DA) around event times and appended them to the trial level dataframe.

Aligned neural signals are appended to the trial level data frames as additional columns. Aligned Z(DA) is baselined by subtracting out its value at “outcome” time (we denote this procedure as de-base for simplicity). For trial average dopamine visualization, we first identified a list of relevant columns as grouping variables of interest, we then dropped any row with one or more NaN entries in the relevant columns. The trial averaged plots are then plotted using seaborn with the error bar being bootstrapped confidence intervals. When reporting summary statistics, we used bootstrapped confidence intervals to characterize mean estimates. For difference comparisons, we used suitable paired or unpaired tests and reported both the testing statistics as well as effect size, and the confidence intervals associated with it.

### Linear Mixed Effects Modeling

Linear Mixed Effect Modeling was used to capture the the effect of past rewards on the current chosen port, or the opposing port. We modeled using rpy2 with the lme4 package with the following formula:

~~~
DA ^∼^ Reward:Switch|Subject + R_chosen:Switch|Subject + Switch|Subject +
     R_unchosen:Switch|Subject + 1|Subject + Reward:Stay|Subject +
     R_chosen:Stay|Subject + R_unchosen:Stay|Subject
~~~

For model simulated behaviors, we changed the output variable to be RPE predictions instead and ran regular regressions without subject level effects due to the uniform sample size in each simulated session.

## QUANTIFICATION AND STATISTICAL ANALYSIS

Statistical analysis was performed using Python with standard packages: scipy, statsmodels, pingouin. We included n=5 subjects with 14 sessions each. Statistical details of each analysis can be found in each figure legend, result section or correspondent method section. Error bars are 95% bootstrapped confidence intervals, and Holm-Bonferroni correction was applied when appropriate.

### Model fitting and comparison

We fit all models to 14 sessions of all 5 mice during the Probswitch phase using custom implemented cogmodels python module. The module finds the best-fitting model parameters for each animal by maximum likelihood estimation. The model initializes the parameters by randomly sampling from distributions as follows:

- RL4p: *α*^+^ *∈* Unif(0, 1), *α*^−^ ∼ Unif(0, 1), *ϕ* ∼ Γ(2, 0.2), *β* ∼ Exp(1)
- RLCF: Same as RL4p
- RFLR: *α* ∼ Exp(1), *ϕ* ∼ *N* (0, 1), *τ* ∼ Exp(1)
- RLFQ3p: *α*^+^ *∈* Unif(0, 1), *α*^−^ ∼ Unif(0, 1), *β* ∼ Exp(1)
- RL_meta: Same as RL4p, plus ζ ∼ Unif(0, 1), *α*_*ν*_ ∼ Unif(0, 1), *ψ* ∼ Unif(0, 1)
- BIfp: *β* ∼ Exp(1), *ϕ* ∼ Γ(2, 0.2), *q* ∼ Unif(0, 0.05)
- BRLfwr: Same parameters as BIfp, plus *α* ∼ Unif(0, 1)

where Γ(*α, θ*) denotes the gamma distribution with shape *α* and scale *β*;𝒩 (*µ, ν*^2^) denotes the Gaussian distribution with mean *µ* and variance *ν*^2^.

After obtaining maximum likelihood estimates, we constructed an empirical range of parameters and performed model recovery tests. We simulated behavioral data from randomly sampled model parameters for each model from the empirical ranges, and refit to the simulated data to obtain the model-fitted values for 1000 random iterations. All three model classes are able to recover model parameters with high correlation (Fig. S2). We performed model comparison by computing the Akaike Information Criterion (AIC) for each model *M*_*i*_:

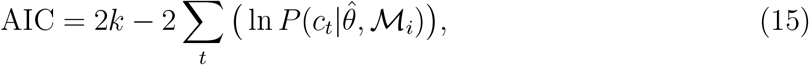

where 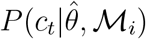 is the likelihood of the data *c*_*t*_ conditional on the maximum likelihood estimate 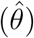. We report AIC scores relative to RL4p, the baseline RL model. More negative scores indicate that a model explains the data better compared to RL4p.

### Model Identification

The model identification tests go through three stages: model behavior simulation, model cross-fitting, and confusion matrix construction. We denote our total model set as *M*.

#### Model behavior simulation

Similar to the previous section, sampling randomly from the empirical range of parameters constructed from the mice data fitting, we simulated behavioral data *D*_*k*_(*m*_*i*_) for 1000 times for each model *m*_*i*_ ∈ *M*, where *k* indexes simulation runs.

#### Model cross-fitting

For each simulated data set *D*_*k*_(*m*_*i*_), we fit each *m*_*j*_ *∈ M*, and get a best fitting model 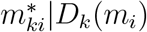 based on lowest AIC measure.

#### Confusion matrix

For all 1000 sessions, we can compute an empirical probability that the model *m*_*j*_ best fits to *D*(*m*_*i*_), or *P* (*m*_*j*_|*m*_*i*_). Then we construct a confusion matrix from *P* (*m*_*j*_|*m*_*i*_), ∀*m*_*i*_, *m*_*j*_ *∈ M*, where each element at row *i*, column *j* represent *P* (*m*_*j*_|*m*_*i*_).

## KEY RESOURCES TABLE

| **REAGENT or RESOURCE** | **SOURCE** | **IDENTIFIER** |
| --- | --- | --- |
| **Antibodies** |  |  |
| 1:1000 chicken anti-GFP | Aves Labs, Inc. | GFP-1020 |
| 1:1000 goat anti-chicken AlexaFluor 488 | Invitrogen by Thermo Fisher Scientific | A11039 |
| **Bacterial and Virus Strains** |  |  |
| AAV5-hSyn-dLight1.2 | Patriarchi et al., 2018; Courtesy of Prof. Lin Tian (UC Davis) | Addgene Cat: 111068-AAV5; RRID: Addgene_111068 |
| AAV9-CAG-dLight1.3b | Patriarchi et al., 2018, Addgene | Addgene Cat: 125560-AAV9; RRID: Addgene_125560 |
| **Chemicals, Peptides, and Recombinant Proteins** |  |  |
| 4% paraformaldehyde (PFA) | This paper | N/A |
| 0.1M phosphate buffer (PB) | This paper | PB |
| Fluoromount-G | Southern Biotech | Cat: 0100-01 |
| **Experimental Models: Organisms/Strains** |  |  |
| C57 Bl/6 male | bred in-house | N/A |
| **Software and Algorithms** |  |  |
| Standard python open source packages (numpy, statsmodels, pingouin, seaborn, plotly, scipy, etc.) | python | pip OR anaconda |
| photometry baseline filtering procedure | Martianova et al., 2019 | DOI:10.3791/60278-v |

## Notes

### Competing Interest Statement

The authors have declared no competing interest.

### Summary of Updates

Figure 4 was revised, and Section on dopamine data updated to included novel data modeling; Supplemental files updated.

## References

[1] Richard S. Sutton and Andrew G. Barto. Reinforcement learning: an introduction. Second edition. Adaptive computation and machine learning series. Cambridge, Massachusetts: The MIT Press, 2018. ISBN: 978-0-262-03924-6.

[2] Jan Gläscher, Alan N. Hampton, and John P. O’Doherty. “Determining a Role for Ventromedial Prefrontal Cortex in Encoding Action-Based Value Signals During Reward-Related Decision Making”. en. In: Cerebral Cortex 19.2 (Feb. 2009), pp. 483–495. ISSN: 1460-2199, 1047-3211. doi: 10.1093/cercor/bhn098. URL: https://academic.oup.com/cercor/article-lookup/doi/10.1093/cercor/bhn098 (visited on 09/27/2023).

[3] Tobias U. Hauser et al. “Cognitive flexibility in adolescence: Neural and behavioral mechanisms of reward prediction error processing in adaptive decision making during development”. en. In: NeuroImage 104 (Jan. 2015), pp. 347–354. ISSN: 10538119. doi: 10.1016/j.neuroimage.2014.09.018. URL: https://linkinghub.elsevier.com/retrieve/pii/S1053811914007605 (visited on 09/27/2023).

[4] Samantha R. Santacruz et al. “Caudate Microstimulation Increases Value of Specific Choices”. en. In: Current Biology 27.21 (Nov. 2017), 3375–3383.e3. ISSN: 09609822. doi: 10.1016/j.cub.2017.09.051. URL: https://linkinghub.elsevier.com/retrieve/pii/S0960982217312496 (visited on 09/27/2023).

[5] Pr Montague, P Dayan, and Tj Sejnowski. “A framework for mesencephalic dopamine systems based on predictive Hebbian learning”. en. In: J. Neurosci. 16.5 (Mar. 1996), pp. 1936–1947. ISSN: 0270-6474, 1529-2401. doi: 10.1523/JNEUROSCI.16-05-01936. 1996. URL: https://www.jneurosci.org/lookup/doi/10.1523/JNEUROSCI.16-05-01936.1996 (visited on 07/30/2023).

[6] Wolfram Schultz, Peter Dayan, and P. Read Montague. “A Neural Substrate of Prediction and Reward”. en. In: Science 275.5306 (Mar. 1997), pp. 1593–1599. ISSN: 0036-8075, 1095-9203. doi: 10.1126/science.275.5306.1593. URL: https://www.science.org/doi/10.1126/science.275.5306.1593 (visited on 04/19/2023).

[7] Neir Eshel et al. “Dopamine neurons share common response function for reward prediction error”. en. In: Nat Neurosci 19.3 (Mar. 2016), pp. 479–486. ISSN: 1097-6256, 1546-1726. doi: 10.1038/nn.4239. URL: https://www.nature.com/articles/nn.4239 (visited on 09/23/2023).

[8] Yuji K. Takahashi et al. “Dopaminergic prediction errors in the ventral tegmental area reflect a multithreaded predictive model”. en. In: Nat Neurosci 26.5 (May 2023), pp. 830–839. ISSN: 1097-6256, 1546-1726. doi: 10.1038/s41593-023-01310-x. URL: https://www.nature.com/articles/s41593-023-01310-x (visited on 08/30/2024).

[9] Benedicte M. Babayan, Naoshige Uchida, and Samuel. J. Gershman. “Belief state representation in the dopamine system”. en. In: Nat Commun 9.1 (May 2018), p. 1891. ISSN: 2041-1723. doi: 10.1038/s41467-018-04397-0. URL: https://www.nature.com/articles/s41467-018-04397-0 (visited on 04/17/2023).

[10] Clara Kwon Starkweather et al. “Dopamine reward prediction errors reflect hidden-state inference across time”. en. In: Nat Neurosci 20.4 (Apr. 2017), pp. 581–589. ISSN: 1097-6256, 1546-1726. doi: 10.1038/nn.4520. URL: https://www.nature.com/articles/nn.4520 (visited on 09/23/2023).

[11] Samuel J. Gershman and Naoshige Uchida. “Believing in dopamine”. en. In: Nat Rev Neurosci 20.11 (Nov. 2019), pp. 703–714. ISSN: 1471-003X, 1471-0048. doi: 10.1038/s41583-019-0220-7. URL: http://www.nature.com/articles/s41583-019-0220-7 (visited on 04/17/2023).

[12] Vincent D. Costa et al. “Reversal Learning and Dopamine: A Bayesian Perspective”. en. In: J. Neurosci. 35.6 (Feb. 2015), pp. 2407–2416. ISSN: 0270-6474, 1529-2401. doi: 10.1523/JNEUROSCI.1989-14.2015. URL: https://www.jneurosci.org/lookup/doi/10.1523/JNEUROSCI.1989-14.2015 (visited on 08/08/2023).

[13] Armin Lak et al. “Midbrain Dopamine Neurons Signal Belief in Choice Accuracy during a Perceptual Decision”. en. In: Current Biology 27.6 (Mar. 2017), pp. 821–832. ISSN: 09609822. doi: 10.1016/j.cub.2017.02.026. URL: https://linkinghub.elsevier.com/retrieve/pii/S096098221730163X (visited on 09/23/2023).

[14] Marta Blanco-Pozo, Thomas Akam, and Mark E. Walton. “Dopamine-independent effect of rewards on choices through hidden-state inference”. en. In: Nat Neurosci (Jan. 2024). ISSN: 1097-6256, 1546-1726. doi: 10.1038/s41593-023-01542-x. URL: https://www.nature.com/articles/s41593-023-01542-x (visited on 03/08/2024).

[15] Jay A Hennig et al. “Emergence of belief-like representations through reinforcement learning”. In: PLOS Computational Biology 19.9 (2023), e1011067.

[16] Lung-Hao Tai et al. “Transient stimulation of distinct subpopulations of striatal neurons mimics changes in action value”. en. In: Nat Neurosci 15.9 (Sept. 2012), pp. 1281–1289. ISSN: 1097-6256, 1546-1726. doi: 10.1038/nn.3188. URL: http://www.nature.com/articles/nn.3188 (visited on 04/18/2023).

[17] Maria K. Eckstein et al. “Reinforcement learning and Bayesian inference provide complementary models for the unique advantage of adolescents in stochastic reversal”. en. In: Developmental Cognitive Neuroscience 55 (June 2022), p. 101106. ISSN: 18789293. doi: 10.1016/j.dcn.2022.101106. URL: https://linkinghub.elsevier.com/retrieve/pii/S1878929322000494 (visited on 04/17/2023).

[18] Celia C. Beron et al. “Mice exhibit stochastic and efficient action switching during probabilistic decision making”. en. In: Proc. Natl. Acad. Sci. U.S.A. 119.15 (Apr. 2022), e2113961119. ISSN: 0027-8424, 1091-6490. doi: 10.1073/pnas.2113961119. URL: https://pnas.org/doi/full/10.1073/pnas.2113961119 (visited on 04/21/2023).

[19] Ethan S. Bromberg-Martin et al. “A Pallidus-Habenula-Dopamine Pathway Signals Inferred Stimulus Values”. en. In: Journal of Neurophysiology 104.2 (Aug. 2010), pp. 1068–1076. ISSN: 0022-3077, 1522-1598. doi: 10.1152/jn.00158.2010. URL: https://www.physiology.org/doi/10.1152/jn.00158.2010 (visited on 06/08/2023).

[20] Cooper D. Grossman, Bilal A. Bari, and Jeremiah Y. Cohen. “Serotonin neurons modulate learning rate through uncertainty”. In: Current Biology 32.3 (2022), 586–599.e7. ISSN: 0960-9822. doi: 10.1016/j.cub.2021.12.006. URL: https://www.sciencedirect.com/science/article/pii/S0960982221016821.

[21] Anne Collins and Etienne Koechlin. “Reasoning, Learning, and Creativity: Frontal Lobe Function and Human Decision-Making”. In: PLOS Biology 10.3 (Mar. 2012), pp. 1–16. doi: 10.1371/journal.pbio.1001293. URL: https://doi.org/10.1371/journal.pbio.1001293.

[22] Milena Rmus, Samuel D McDougle, and Anne Ge Collins. “The role of executive function in shaping reinforcement learning”. en. In: Current Opinion in Behavioral Sciences 38 (Apr. 2021), pp. 66–73. ISSN: 23521546. doi: 10.1016/j.cobeha.2020.10.003. URL: https://linkinghub.elsevier.com/retrieve/pii/S2352154620301480 (visited on 09/26/2023).

[23] HyungGoo R. Kim et al. “A Unified Framework for Dopamine Signals across Timescales”. en. In: Cell 183.6 (Dec. 2020), 1600–1616.e25. ISSN: 00928674. doi: 10.1016/j.cell.2020.11.013. URL: https://linkinghub.elsevier.com/retrieve/pii/S0092867420315300 (visited on 09/23/2023).

[24] KevinT. Beier et al. “Circuit Architecture of VTA Dopamine Neurons Revealed by Systematic Input-Output Mapping”. In: Cell 162.3 (2015), pp. 622–634. ISSN: 0092-8674. doi: 10.1016/j.cell.2015.07.015. URL: https://www.sciencedirect.com/science/article/pii/S0092867415008521.

[25] Ronald Keiflin and Patricia H. Janak. “Dopamine Prediction Errors in Reward Learning and Addiction: From Theory to Neural Circuitry”. In: Neuron 88.2 (2015), pp. 247–263. ISSN: 0896-6273. doi: 10.1016/j.neuron.2015.08.037. URL: https://www.sciencedirect.com/science/article/pii/S089662731500731X.

[26] Matthijs AA van der Meer and A David Redish. “Ventral striatum: a critical look at models of learning and evaluation”. en. In: Current Opinion in Neurobiology 21.3 (June 2011), pp. 387–392. ISSN: 09594388. doi: 10.1016/j.conb.2011.02.011. URL: https://linkinghub.elsevier.com/retrieve/pii/S0959438811000389 (visited on 04/18/2023).

[27] Tommaso Patriarchi et al. “Ultrafast neuronal imaging of dopamine dynamics with designed genetically encoded sensors”. In: Science 360.6396 (2018), eaat4422. doi: 10.1126/science.aat4422. eprint: https://www.science.org/doi/pdf/10.1126/science.aat4422. URL: https://www.science.org/doi/abs/10.1126/science.aat4422.

[28] Makoto Ito and Kenji Doya. “Validation of decision-making models and analysis of decision variables in the rat basal ganglia”. In: Journal of Neuroscience 29.31 (2009), pp. 9861–9874.

[29] Maria K. Eckstein and Anne G. E. Collins. “Computational evidence for hierarchically structured reinforcement learning in humans”. en. In: Proc. Natl. Acad. Sci. U.S.A. 117.47 (Nov. 2020), pp. 29381–29389. ISSN: 0027-8424, 1091-6490. doi: 10.1073/pnas.1912330117. URL: https://pnas.org/doi/full/10.1073/pnas.1912330117 (visited on 09/26/2023).

[30] Eldad Yechiam et al. “Using cognitive models to map relations between neuropsychological disorders and human decision-making deficits”. In: Psychological science 16.12 (2005), pp. 973–978.

[31] Michael J Frank et al. “Genetic triple dissociation reveals multiple roles for dopamine in reinforcement learning”. In: Proceedings of the National Academy of Sciences 104.41 (2007), pp. 16311–16316.

[32] Yael Niv et al. “Neural prediction errors reveal a risk-sensitive reinforcement-learning process in the human brain”. In: Journal of Neuroscience 32.2 (2012), pp. 551–562.

[33] Samuel J Gershman. “Do learning rates adapt to the distribution of rewards?” In: Psychonomic Bulletin & Review 22 (2015), pp. 1320–1327.

[34] Michiyo Sugawara and Kentaro Katahira. “Dissociation between asymmetric value updating and perseverance in human reinforcement learning”. In: Scientific Reports 11.1 (2021), p. 3574.

[35] Erie D Boorman, Timothy E Behrens, and Matthew F Rushworth. “Counterfactual choice and learning in a neural network centered on human lateral frontopolar cortex”. In: PLoS Biology 9.6 (2011), e1001093.

[36] Stefano Palminteri et al. “The computational development of reinforcement learning during adolescence”. In: PLoS Computational Biology 12.6 (2016), e1004953.

[37] Stefano Palminteri et al. “Confirmation bias in human reinforcement learning: Evidence from counterfactual feedback processing”. In: PLoS computational biology 13.8 (2017), e1005684.

[38] Alan N Hampton, Peter Bossaerts, and John P O’doherty. “The role of the ventromedial prefrontal cortex in abstract state-based inference during decision making in humans”. In: Journal of Neuroscience 26.32 (2006), pp. 8360–8367.

[39] Hannah M. Bayer and Paul W. Glimcher. “Midbrain Dopamine Neurons Encode a Quantitative Reward Prediction Error Signal”. In: Neuron 47.1 (2005), pp. 129–141. ISSN: 0896-6273. doi: 10.1016/j.neuron.2005.05.020. URL: https://www.sciencedirect.com/science/article/pii/S0896627305004678.

[40] Clara Kwon Starkweather and Naoshige Uchida. “Dopamine signals as temporal difference errors: recent advances”. en. In: Current Opinion in Neurobiology 67 (Apr. 2021), pp. 95–105. ISSN: 09594388. doi: 10.1016/j.conb.2020.08.014. URL: https://linkinghub.elsevier.com/retrieve/pii/S095943882030132X (visited on 04/19/2023).

[41] Karyna Mishchanchuk et al. “Hidden state inference requires abstract contextual representations in ventral hippocampus”. In: bioRxiv (2024). doi: 10.1101/2024.05.17.594673. eprint: https://www.biorxiv.org/content/early/2024/05/19/2024.05.17.594673.full.pdf. URL: https://www.biorxiv.org/content/early/2024/05/19/2024.05.17.594673.

[42] NathanielD. Daw et al. “Model-Based Influences on Humans’ Choices and Striatal Prediction Errors”. In: Neuron 69.6 (2011), pp. 1204–1215. ISSN: 0896-6273. doi: 10.1016/j.neuron.2011.02.027. URL: https://www.sciencedirect.com/science/article/pii/S0896627311001255.

[43] Anne G.E. Collins et al. “Working Memory Load Strengthens Reward Prediction Errors”. en. In: J. Neurosci. 37.16 (Apr. 2017), pp. 4332–4342. ISSN: 0270-6474, 1529-2401. doi: 10.1523/JNEUROSCI.2700-16.2017. URL: https://www.jneurosci.org/lookup/doi/10.1523/JNEUROSCI.2700-16.2017 (visited on 09/26/2023).

[44] Anne G.E. Collins. “Learning Structures Through Reinforcement”. en. In: Goal-Directed Decision Making. Elsevier, 2018, pp. 105–123. ISBN: 978-0-12-812098-9. doi: 10.1016/B978-0-12-812098-9.00005-X. URL: https://linkinghub.elsevier.com/retrieve/pii/B978012812098900005X (visited on 09/27/2023).

[45] Aspen H. Yoo, Haley Keglovits, and Anne G. E. Collins. “Lowered inter-stimulus discriminability hurts incremental contributions to learning”. en. In: Cogn Affect Behav Neurosci (Sept. 2023). ISSN: 1530-7026, 1531-135X. doi: 10.3758/s13415-023-01104-5. URL: https://link.springer.com/10.3758/s13415-023-01104-5 (visited on 09/26/2023).

[46] Maël Donoso, Anne G. E. Collins, and Etienne Koechlin. “Foundations of human reasoning in the prefrontal cortex”. en. In: Science 344.6191 (June 2014), pp. 1481–1486. ISSN: 0036-8075, 1095-9203. doi: 10.1126/science.1252254. URL: https://www.science.org/doi/10.1126/science.1252254 (visited on 09/26/2023).

[47] James B. Heald, Máté Lengyel, and Daniel M. Wolpert. “Contextual inference underlies the learning of sensorimotor repertoires”. en. In: Nature 600.7889 (Dec. 2021), pp. 489–493. ISSN: 0028-0836, 1476-4687. doi: 10.1038/s41586-021-04129-3. URL: https://www.nature.com/articles/s41586-021-04129-3 (visited on 09/26/2023).

[48] Anne G. E. Collins and Michael J. Frank. “Cognitive control over learning: Creating, clustering, and generalizing task-set structure.” en. In: Psychological Review 120.1 (Jan. 2013), pp. 190–229. ISSN: 1939-1471, 0033-295X. doi: 10.1037/a0030852. URL: http://doi.apa.org/getdoi.cfm?doi=10.1037/a0030852 (visited on 09/26/2023).

[49] Anne G. E. Collins and Michael J. Frank. “Within- and across-trial dynamics of human EEG reveal cooperative interplay between reinforcement learning and working memory”. en. In: Proc. Natl. Acad. Sci. U.S.A. 115.10 (Mar. 2018), pp. 2502–2507. ISSN: 0027-8424, 1091-6490. doi: 10.1073/pnas.1720963115. URL: https://pnas.org/doi/full/10.1073/pnas.1720963115 (visited on 09/26/2023).

[50] Mark W. Howe et al. “Prolonged dopamine signalling in striatum signals proximity and value of distant rewards”. en. In: Nature 500.7464 (Aug. 2013), pp. 575–579. ISSN: 0028-0836, 1476-4687. doi: 10.1038/nature12475. URL: http://www.nature.com/articles/nature12475 (visited on 04/22/2023).

[51] Karolina Farrell, Armin Lak, and Aman B. Saleem. “Midbrain dopamine neurons signal phasic and ramping reward prediction error during goal-directed navigation”. en. In: Cell Reports 41.2 (Oct. 2022), p. 111470. ISSN: 22111247. doi: 10.1016/j.celrep.2022.111470. URL: https://linkinghub.elsevier.com/retrieve/pii/S2211124722013201 (visited on 08/30/2024).

[52] Maite Azcorra et al. “Unique functional responses differentially map onto genetic subtypes of dopamine neurons”. en. In: Nat Neurosci 26.10 (Oct. 2023), pp. 1762–1774. ISSN: 1097-6256, 1546-1726. doi: 10.1038/s41593-023-01401-9. URL: https://www.nature.com/articles/s41593-023-01401-9 (visited on 06/20/2024).

[53] Munir Gunes Kutlu et al. “Dopamine release in the nucleus accumbens core signals perceived saliency”. en. In: Current Biology 31.21 (Nov. 2021), 4748–4761.e8. ISSN: 09609822. doi: 10.1016/j.cub.2021.08.052. URL: https://linkinghub.elsevier.com/retrieve/pii/S096098222101188X (visited on 06/21/2024).

[54] Munir Gunes Kutlu et al. “Dopamine signaling in the nucleus accumbens core mediates latent inhibition”. en. In: Nat Neurosci 25.8 (Aug. 2022), pp. 1071–1081. ISSN: 1097-6256, 1546-1726. doi: 10.1038/s41593-022-01126-1. URL: https://www.nature.com/articles/s41593-022-01126-1 (visited on 04/22/2023).

[55] Anne L. Collins and Benjamin T. Saunders. “Heterogeneity in striatal dopamine circuits: Form and function in dynamic reward seeking”. In: Journal of Neuroscience Research 98.6 (2020), pp. 1046–1069. doi: 10.1002/jnr.24587. eprint: https://onlinelibrary.wiley.com/doi/pdf/10.1002/jnr.24587. URL: https://onlinelibrary.wiley.com/doi/abs/10.1002/jnr.24587.

[56] Masakazu Taira et al. “Dopamine Release in the Nucleus Accumbens Core Encodes the General Excitatory Components of Learning”. In: Journal of Neuroscience 44.35 (2024). ISSN: 0270-6474. doi: 10.1523/JNEUROSCI.0120-24.2024. eprint: https://www.jneurosci.org/content/44/35/e0120242024.full.pdf. URL: https://www.jneurosci.org/content/44/35/e0120242024.

[57] Yuhao Wang et al. “Dopamine encoding of novelty facilitates efficient uncertainty-driven exploration”. In: PLOS Computational Biology 20.4 (Apr. 2024), pp. 1–27. doi: 10.1371/journal.pcbi.1011516. URL: https://doi.org/10.1371/journal.pcbi.1011516.

[58] Luke T. Coddington, Sarah E. Lindo, and Joshua T. Dudman. “Mesolimbic dopamine adapts the rate of learning from action”. en. In: Nature 614.7947 (Feb. 2023), pp. 294–302. ISSN: 0028-0836, 1476-4687. doi: 10.1038/s41586-022-05614-z. URL: https://www.nature.com/articles/s41586-022-05614-z (visited on 10/24/2023).

[59] Huijeong Jeong et al. “Mesolimbic dopamine release conveys causal associations”. en. In: Science 378.6626 (Dec. 2022), eabq6740. ISSN: 0036-8075, 1095-9203. doi: 10.1126/science.abq6740. URL: https://www.science.org/doi/10.1126/science.abq6740 (visited on 06/21/2024).

[60] Thomas Akam, Rui Costa, and Peter Dayan. “Simple Plans or Sophisticated Habits? State, Transition and Learning Interactions in the Two-Step Task”. In: PLOS Computational Biology 11.12 (Dec. 2015), pp. 1–25. doi: 10.1371/journal.pcbi.1004648. URL: https://doi.org/10.1371/journal.pcbi.1004648.

[61] Kevin J Miller, Matthew M Botvinick, and Carlos D Brody. “Dorsal hippocampus contributes to model-based planning”. en. In: Nat Neurosci 20.9 (Sept. 2017), pp. 1269–1276. ISSN: 1097-6256, 1546-1726. doi: 10.1038/nn.4613. URL: https://www.nature.com/articles/nn.4613 (visited on 11/08/2023).

[62] Ekaterina Martianova, Sage Aronson, and Christophe D. Proulx. “Multi-Fiber Photometry to Record Neural Activity in Freely-Moving Animals”. In: JoVE 152 (Oct. 2019), e60278. ISSN: 1940-087X. doi: 10.3791/60278. URL: https://doi.org/10.3791/60278.

[63] Kenji Doya et al. “Multiple model-based reinforcement learning”. In: Neural computation 14.6 (2002), pp. 1347–1369.

