## Supplementary Figures for "Nucleus accumbens dopamine release reflects Bayesian inference during instrumental learning"

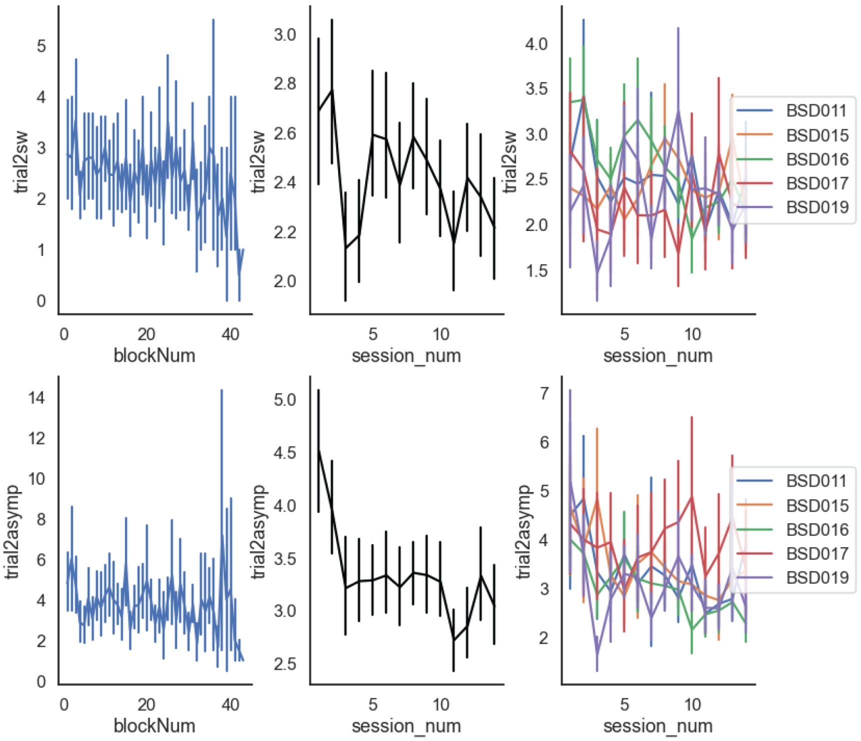


Figure S1: **Mice switch and commit to correct ports faster across multiple sessions.** Top row shows the results for trial2sw, defined as the number of trials taken for mice to switch to the correct port in a new block. Bottom row shows the results for trial2asymp, defined as the first trial from which the animal chose the correct port and then persisted with the choice selection for 2 subsequent trials consequently. First column shows the relationship between the measures and the number of blocks within a training session, demonstrating within session learning. A downward trend for both measures suggests learning and faster switches across different reward blocks within session. Second column shows that the switch measures decreased as the number of training sessions increased, describing learning across multiple sessions. This is consistent with results in Fig. 1F, where the correct rate of the mice improved over multiple sessions. Third column shows the same thing as the second column, but separated by animals.

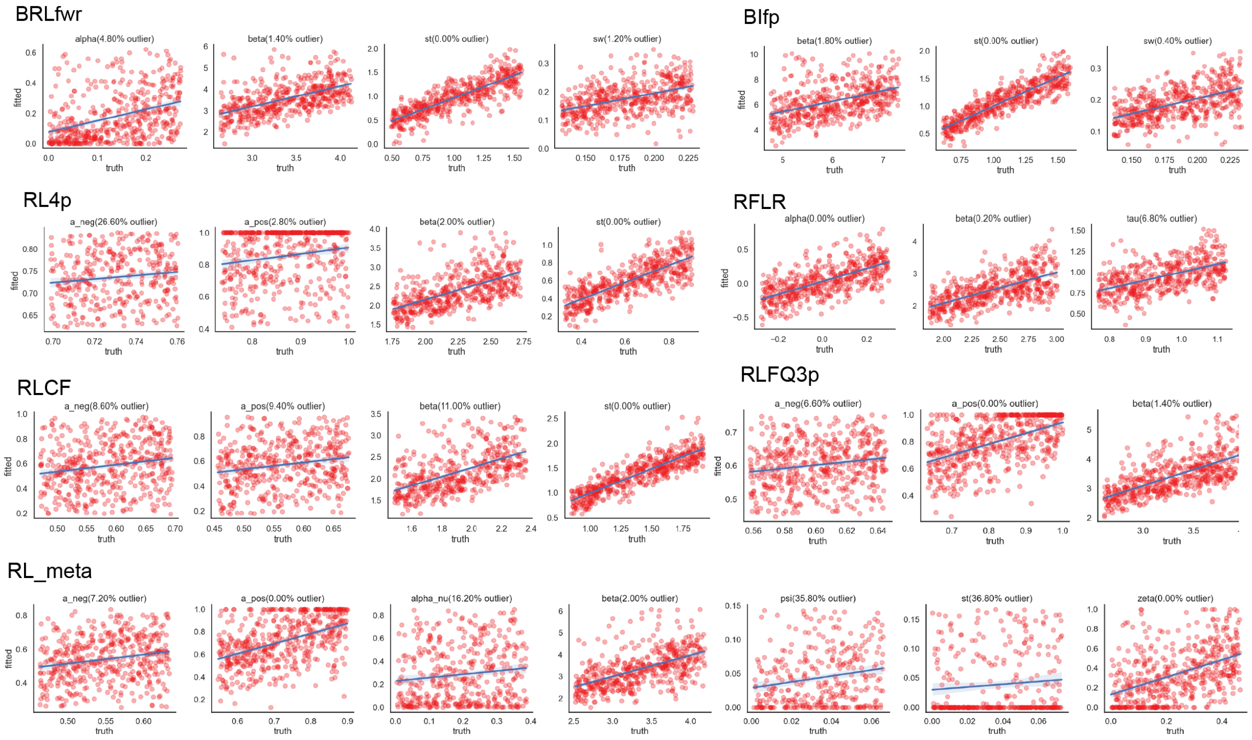


Figure S2: **Parameter recovery of cognitive models.** We took the maxima and minima for each of the best fitted parameter values for all subjects, and constructed a uniform distribution for each. We then sampled from these empirical distributions for simulating behaviors and fitted best fitted parameters for each set of simulated behaviors for each model. The results are generated after 500 runs of random parameter samples for each model, shown as a scatter plot with truth parameter against fitted parameters. Each row is a different model noted at the top left corner. Outliers, defined as 6 standard deviations from mean true parameter values, were thrown out for visualization purposes but percentages were noted.

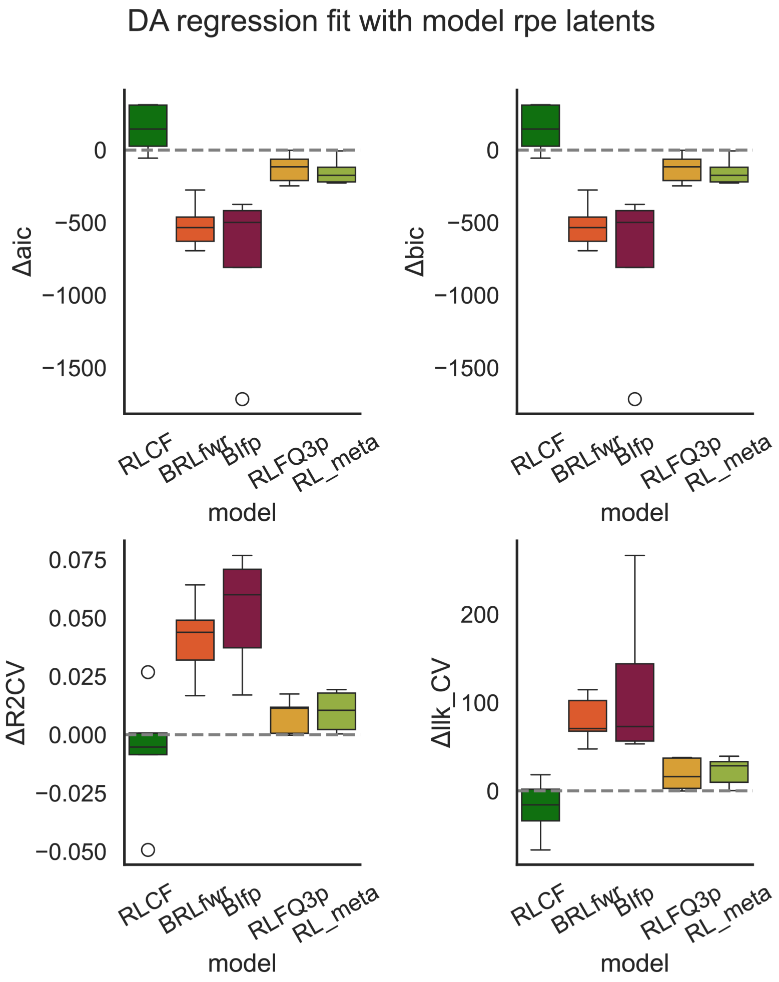


Figure S3: **Comparison of fitness of model predicted RPE to dopamine data.** Similar to Fig. 4F, we included results of model fitness using different metrics: relative AIC, relative BIC, relative cross validated R2 (R2CV), and relative cross validated log likelihood (llk_CV). All metrics converged on favoring Bayesian model predictions of dopamine at outcome phase.

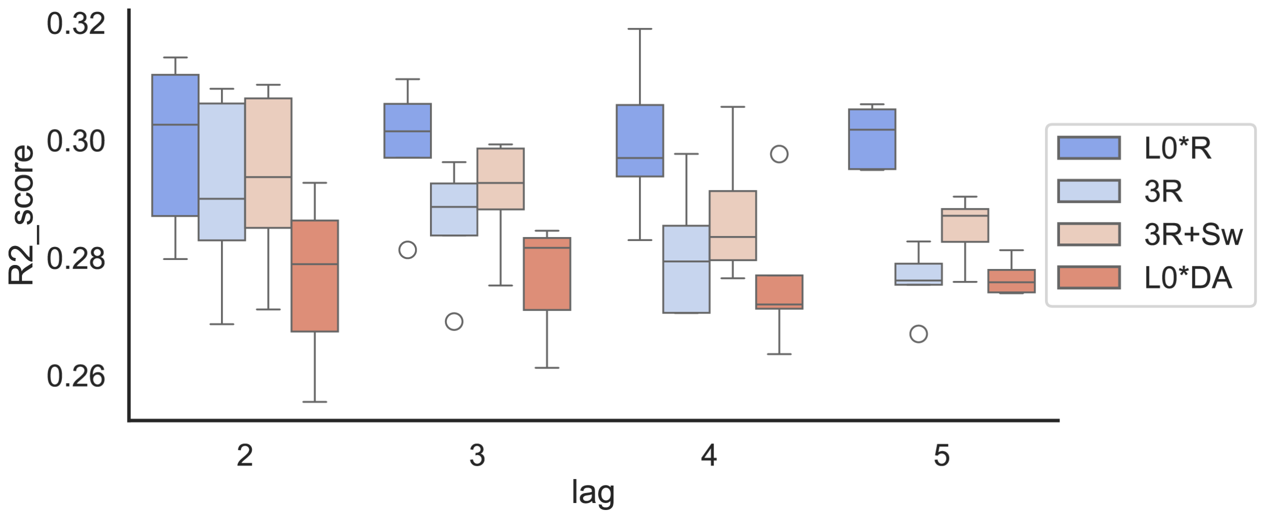


Figure S4: **Selection of LMER models and lags.** We compared the cross validated R2 score using different feature sets, while keeping the use of dopamine as output variable. L0:R: Standard formulation using N trial back choice and reward interactions. 3R: just R_chosen, R_unchosen, Reward features. 3R+Sw: in addition to 3R features, we included interactions of whether a trial is an animal switch trial, or animal stay trials. L0:DA: we used the interactions between past trial dopamine values and choice selections. Together we found that 3R+Sw was enough to capture similar levels of variance compared to L0:R. We used 4 lags because it allows us to capture a relatively high amount of past outcome rewards levels, with only the expenses of 1% variance explained.

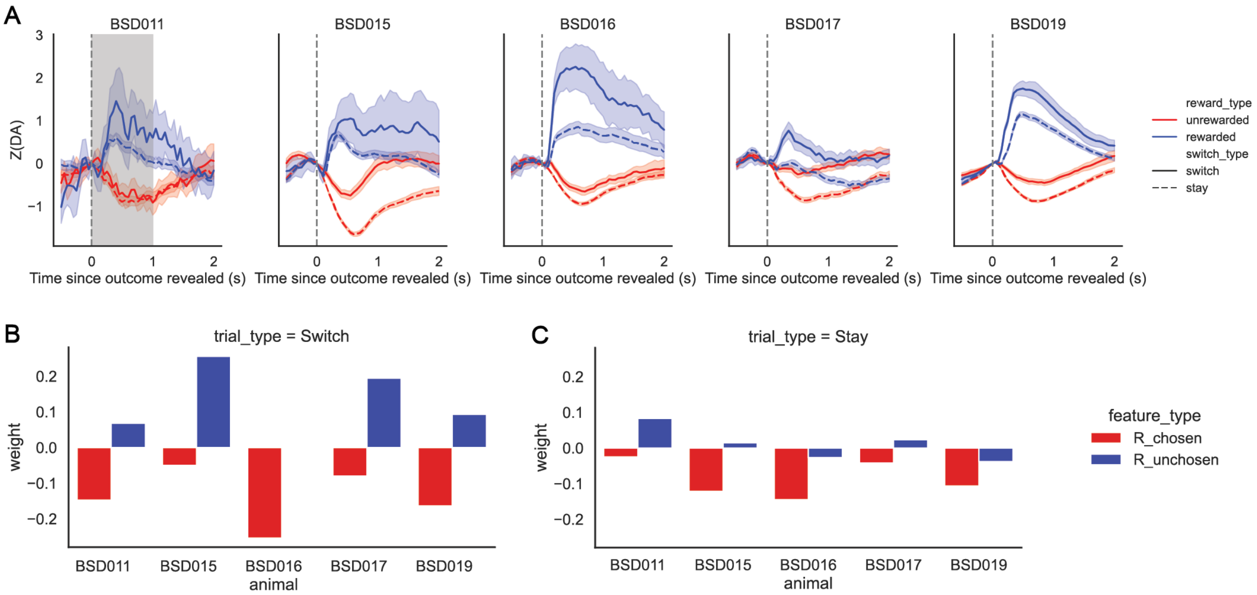


Figure S5: **Dopamine data are qualitatively consistent across animals.** (A) Average differences in dopamine responses to rewards or unrewarded outcomes across switch stay trials are fairly consistent across animals. (B-C) Similar to LMER regression of influence of past rewards on dopamine responses, we did OLS for each animal separately. All but one animal shows a strong qualitative resemblance to Bayesian model predictions for Switch trials. As discussed in the main text, R_unchosen effect is highly variable across animals, and has a near-zero net effect, corresponding more to the predictions of BIfp simulation.
